# A mouse tissue atlas of small non-coding RNA

**DOI:** 10.1101/430561

**Authors:** Alina Isakova, Tobias Fehlmann, Andreas Keller, Stephen R. Quake

**Affiliations:** Department of Bioengineering, Stanford University, Stanford, California, USA; Chair for Clinical Bioinformatics, Saarland University, 66123 Saarbrücken, Germany; School of Medicine Office, Stanford University, Stanford, CA 94305, USA; Department of Applied Physics, Stanford University, Stanford, California, USA; Chan Zuckerberg Biohub, San Francisco, California, USA

## Abstract

Small non-coding RNAs (ncRNAs) play a vital role in a broad range of biological processes both in health and disease. A comprehensive quantitative reference of small ncRNA expression would significantly advance our understanding of ncRNA roles in shaping tissue functions. Here, we systematically profiled the levels of five ncRNA classes (miRNA, snoRNA, snRNA, scaRNA and tRNA fragments) across eleven mouse tissues by deep sequencing. Using fourteen biological replicates spanning both sexes, we identified that ~ 30% of small ncRNAs are distributed across the body in a tissue-specific manner with some are also being sexually dimorphic. We found that miRNAs are subject to “arm switching” between healthy tissues and that tRNA fragments are retained within tissues in both a gene- and a tissue-specific manner. Out of eleven profiled tissues we confirmed that brain contains the largest number of unique small ncRNA transcripts, some of which were previously annotated while others are identified for the first time in this study. Furthermore, by combining these findings with single-cell ATAC-seq data, we were able to connect identified brain-specific ncRNA with their cell types of origin. These results yield the most comprehensive characterization of specific and ubiquitous small RNAs in individual murine tissues to date, and we expect that this data will be a resource for the further identification of ncRNAs involved in tissue-function in health and dysfunction in disease.

**HIGHLIGHTS:**

- An atlas of tissue levels of multiple small ncRNA classes generated from 14 biological replicates of both sexes across 11 tissues
- Distinct distribution patterns of miRNA arms and tRNA fragments across tissues suggest the existence of tissue-specific mechanisms of ncRNA cleavage and retention
- miRNA expression is sex specific in healthy tissues
- Small RNA-seq and scATAC-seq data integration produce a detailed map of cell-type specific ncRNA profiles in the mouse brain

## INTRODUCTION

Small non-coding RNAs (ncRNAs) are a large family of endogenously expressed transcripts, 18 to 200 nucleotides long, that play a crucial role in regulating cell function (Bartel, 2018; Cech and Steitz, 2014). Seen mainly as “junk” RNA of unknown function two decades ago, today small ncRNAs are believed to be involved in nearly all developmental and pathological processes in mammals (Cech and Steitz, 2014; Esteller, 2011; Gebetsberger et al., 2017; He and Hannon, 2004; Ng et al., 2016). While the exact function of many ncRNAs remain unknown, numerous studies have revealed the direct involvement of various small ncRNAs in regulation of gene expression at the levels of post-transcriptional mRNA processing (Becker et al., 2019; Gebert and MacRae, 2019; Kuscu et al., 2018; Telonis et al., 2015a) and ribosome biogenesis (Reichow et al., 2007). Aberrant expression of small ncRNAs, in turn, has been associated with diseases such as cancer, autoimmune disease and several neurodegenerative disorders (Almeida et al., 2011; Keller et al., 2011; Liu et al., 2017; Somel et al., 2010).

Mammalian cells express several classes of small ncRNA including microRNA (miRNA) (Ha and Kim, 2014), small interfering RNAs (siRNA), small nucleolar RNAs (snoRNA) (Matera et al., 2007), small nuclear RNA (snRNA)(Kiss, 2004), PIWI-interacting RNA (piRNA) (Ishizu et al., 2012), tRNA-derived small RNAs (tRFs) (Kumar et al., 2014), with some being shown to be expressed in a tissue- (Dittmar et al., 2006; Landgraf et al., 2007), cell-type-(Faridani et al., 2016; Volinia et al., 2006) or even cell-state-specific manner (Hayes et al., 2014; Palfi et al., 2016; Sherstyuk et al., 2017). Through their interactions with ribosomes and mRNA that code for protein expression, these small non-coding molecules shape the dynamic molecular spectrum of tissues (Faridani et al., 2016; Gebetsberger et al., 2017; Kuscu et al., 2018). Despite extensive knowledge of ncRNA biogenesis and function (Bartel, 2018; Jorjani et al., 2016; Kumar et al., 2016), much remains to be explored about tissue- and sex-specific small ncRNA expression. Given the emerging role of ncRNAs as biomarkers (Anfossi et al., 2018; Slack and Chinnaiyan, 2019) and potent therapeutic targets (Janssen et al., 2013), a comprehensive reference atlas of tissue small ncRNA expression would represent a valuable resource not only for fundamental but also clinical research.

The first attempts to catalog tissue-specific mammalian small ncRNAs began a decade ago with characterization of miRNA levels (Ach et al., 2008; Hsu et al., 2007; Landgraf et al., 2007; Liang et al., 2007). While these pioneering microarray-, qPCR- and Sanger sequencing-based studies mapped only a limited number of highly expressed miRNA, they nevertheless established a “gold standard” reference for the following decade of miRNA research. Efforts to characterize tissue-specific non-coding transcripts have recently resumed with employment of RNA-seq, which greatly advanced the discovery of novel and previously undetected miRNA (Londin et al., 2015a; McCall et al., 2017; de Rie et al., 2017). However, none of the prior studies encompass a spectrum of mammalian tissues from both female and male individuals, let alone the other non-coding RNA types that were recently identified to carry-out tissue- and cell-type specific functions (Jehn et al., 2019; Kuscu et al., 2018; Rimer et al., 2018).

Here, we describe a comprehensive atlas of small ncRNA expression across eleven mammalian tissues. Using multiple biological replicates (n=14) from individuals of both sexes, we mapped tissue specific as well as broadly transcribed small ncRNA attributed to five different classes and spanning a large spectrum of expression levels. Our data reveals that tissue specificity extends to ncRNA types other than miRNA and provides novel insights on tissue-dependent distribution of miRNA arms and tRNA fragments. We have also discovered that certain miRNAs are broadly sexually dimorphic while other show sex-bias in the context of specific tissues. Finally, integrating our ncRNA expression measurements with the scATAC-seq data (Cusanovich et al., 2018, 10X Genomics) enabled us to map cell-type specificity of small ncRNA expressed in the adult mouse brain.

## RESULTS

### Small ncRNA expression atlas of mouse tissues

We profiled the expression of small ncRNA across ten tissues from adult female (n=10) and eleven from adult male (n=4) C57BL/6J mice (**Figure 1A, Table S1**). We generated a dataset comprising in total of 140 small ncRNA sequencing libraries from brain, lung, heart, muscle, kidney, pancreas, liver, small intestine, spleen, bone marrow and testes RNA. Each library yielded ~5-20 million reads mapping to the mouse genome, out of which, on average ~ 7 million mapped to the exons of small ncRNA genes (see ***Methods***), resulting in the total of ~ 100 million ncRNA reads per tissue (**Figure S1A**). Using the GENCODE M20 (Frankish et al., 2019), GtRNAdb (Chan and Lowe, 2016) and miRBase (Kozomara and Griffiths-Jones, 2014) mouse annotations, we mapped the expression of distinct small ncRNA classes: miRNA, snRNA, snoRNA, scaRNA, tRF and other small ncRNA in profiled tissues (**Figure 1B**). Among all the tissues we identified a total of 1317 distinct miRNA, 733 snRNA, 583 snoRNA, 25 scaRNA, 346 tRNA, 22 mitochondrial tRNA and 193 other miscellaneous small ncRNA genes, which corresponds to 60%, 53%, 39% 96%, 92%, 100% and 34% of annotated transcripts of each respective class (**Figure 1B**). miRNA was the most abundant small ncRNA type in our libraries, followed by snoRNA, snRNA and tRFs (**Figure 1C and Figure S1B**). With respect to protein coding genes, the detected tRFs were intergenic, snoRNA were of intronic origin, snRNA and scaRNA were intronic and intergenic (63/35% and 64/36% respectively) and miRNA were transcribed from either introns (53%), exons (11%) or intergenic regions (11%) (**Figure S1C**). The number of distinct ncRNA greatly varied across tissues; for example lymphoid tissues (lung, spleen and bone marrow) contained the largest number of distinct ncRNA while pancreas and liver contained the lowest (**Figure 1C and Figure S1B**). Furthermore, within the profiled tissues we detected 95.1% of miRNA precursors denoted by miRBase v22 database as high confidence transcripts (Kozomara and Griffiths-Jones, 2014). Using the obtained data, we have reconstructed the genome-wide expression map of various small ncRNA types across eleven murine tissues (**Figure 1D, Table S2, Table 1**).

**Figure 1.**
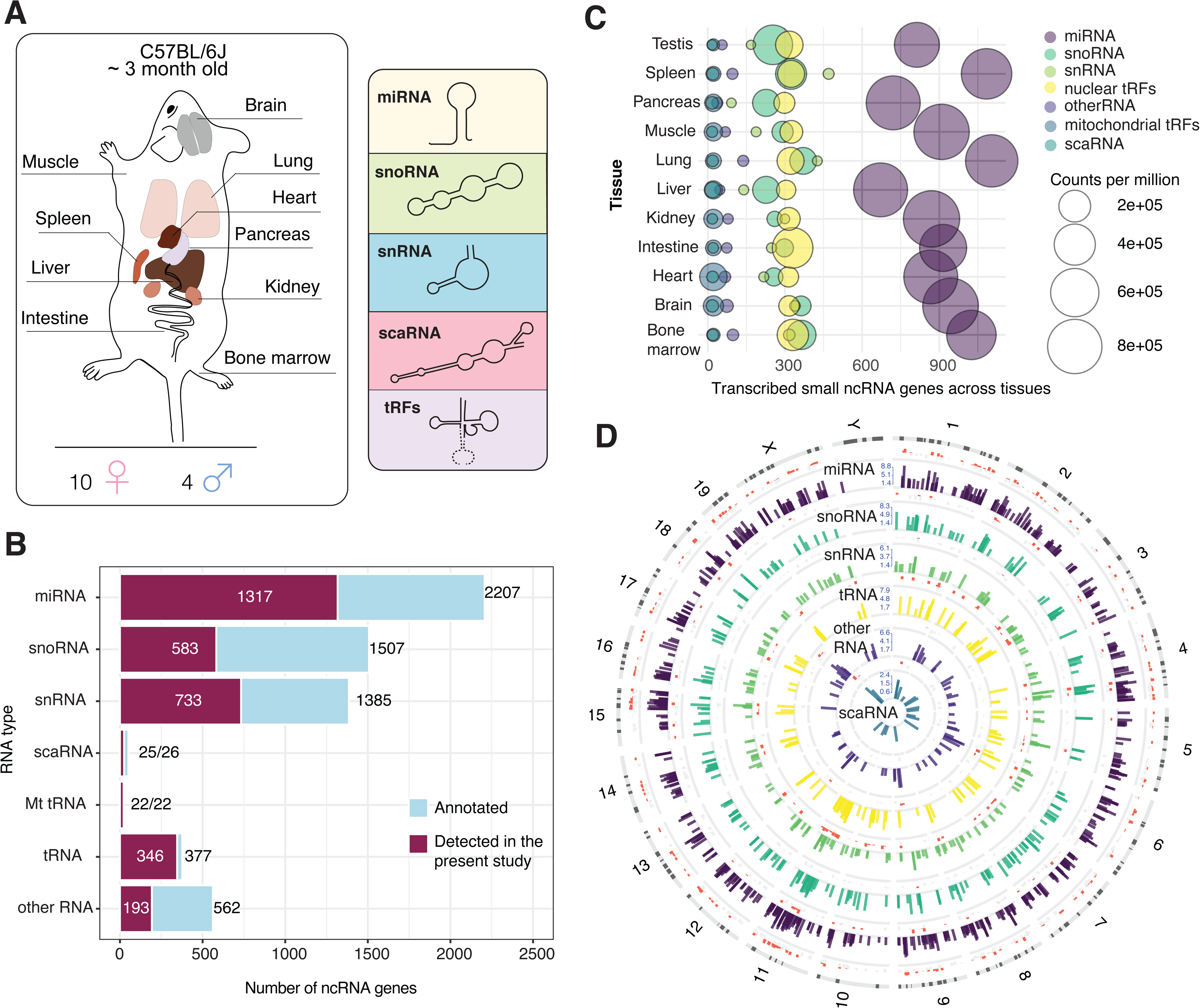
Small ncRNA expression across mouse tissues. **(A) Tissues and ncRNA classes profiled in the current study.** Ten somatic tissues were collected from adult mice (n=14). Testes were collected from male individuals (n=4). **(B) ncRNAs identified in the current study.** Numbers indicate detected and total annotated within GENCODE M20 miRNA, snoRNA, snRNA, scaRNA, Mt_tRNA as well as high-confidence tRNA listed in GtRNAdb. **(C) Coverage of ncRNA types within the profiled tissues.** ncRNA was considered transcribed in a tissue if detected at > 1 cpm. **(D) Genomic map of sRNAs expression across mouse genome.** Bars show the log-transformed normalized expression count of ncRNAs. Red and grey bars around each circle represent the variance of each sRNA across ten mouse tissues. Red denotes highly (the standard deviation of expression above 25% of the mean value) and grey - low variable ncRNAs (standard deviation below 25% of the mean value).

**Table 1.** Studies cataloging small non-coding RNA across tissues using high-through sequencing.

| *Study* | **Organism** | **Total number of small non-coding RNA reads** | **Analyzed small ncRNA types** |
| --- | --- | --- | --- |
| *Landgraf et al. Cell, 2007* | Human | ~300,000 | miRNA |
| *Minami et al., Sci data, 2014* | Rat | ~50,000,000 | miRNA |
| *De Rie et al., Nat Biotechnol., 2017* | Human | ~1,200,000,000 | miRNA |
|  | Mouse | ~ 360,000,000 | miRNA |
| *McCall et al., Genome Res, 2017* | Human | ~2,400,000,000<br>(includes all available public miRNA-seq data) | miRNA |
| *Minami et al., Sci data, 2014* | Rat | ~50,000,000 | miRNA |
| *Sun et al., Sci data, 2019* | Cow | ~103,000,000 | miRNA |
| ***Present study*** | Mouse | ~1,100,000,000 | miRNA, snoRNA, snRNA, scaRNA, miscRNA, tRFs |

### Tissue-specific expression of small ncRNA

We first assessed the differences in the levels of small ncRNAs across profiled tissues at the gene level, based on the expression of all assayed RNA types. Dimensionality reduction via t-distributed stochastic neighbor embedding (tSNE) (Laurens Van Der Maaten and Geoffrey Hinton, 2008) (***Methods***) on ncRNA genes revealed a robust clustering of samples according to tissue types (**Figure 2A**). For each ncRNA we have computed the tissue specificity index, *TSI*, as described previously in (Ludwig et al., 2016). We observed that ~ 17% of all detected ncRNA were present in only one tissue (*TSI* = 1) (**Figure S2A**), while the remaining ncRNA were either ubiquitously expressed or found in some but absent in other tissues. We have next ran a differential gene expression (DGE) analysis on all detected ncRNA across eleven tissues (see ***Methods***) and found that out of 3219 detected genes, 897 (28%) contribute to the tissue-specific signature of ncRNA expression (BH-adjusted p-value < 0.01) (**Figure 2B-C, Table S3**). Interestingly, we found brain to contain the highest number of unique transcripts not present in other tissues (~ 400) (**Figure 2B-C, Table S3**) even though lung, spleen and bone marrow expressed the widest spectrum of detected genes **(Figure 1C)**. We found miRNA to be the main contributor of tissue-specificity (reflected by the lowest p-values in our DGE test) **(Figure 2B-C);** however, we have also identified hundreds of ncRNAs of other types which are expressed in a tissue-specific fashion **(Figure S2)**.

**Figure 2.**
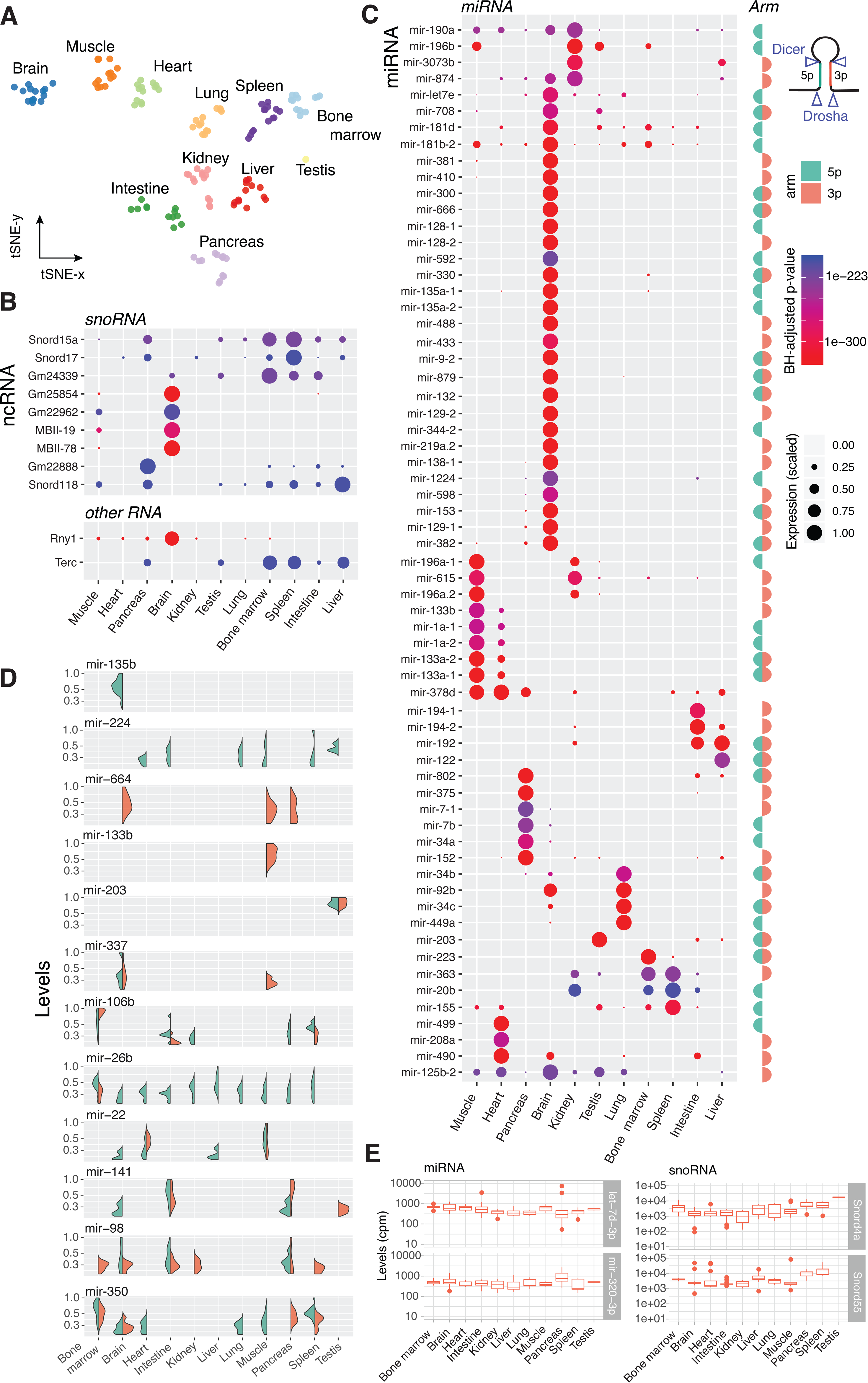
Tissue-specific patterns of small ncRNA expression. **(A) t-SNE projection of ncRNA expression patterns** performed on ~4000 ncRNA genes of various classes detected in eleven mouse tissues. **(B) Dotplot** of tissue specific snoRNA, *Rny1* and *Terc*, identified as the most tissue-specific (BH-adjusted p-value < 1e-150 in LRT test). The size of the dot represents scaled log-transformed normalized counts. **(C) Dotplot** of tissue specific miRNA. Only a subset of passing the specificity threshold (BH-adjusted p-value < 1e-0.01 in LRT test) miRNA is shown. “Arm” column denotes whether *5p-*, *3p-* or both arms are passing the specificity threshold. **(D) Levels of** miRNAs arms detected across tissues. Representative miRNAs, for which we consistently detect ether one of the arms, both or switched arm between tissues. Y-axis represents normalized scaled counts. **(E) Examples** of ubiquitous ncRNA present in all tissues.

### Tissue-specific snoRNAs

We found that snoRNA alone is capable to separate the majority of profiled tissues based on their transcript levels (**Figure S2B**), with over 200 snoRNA showing tissues-specific patterns (**Figure S2C, Table S3)**. For example, we discovered that maternally imprinted *AF357428* (also known as MBII-78), *AF357341* (MBII-19) and *Gm25854* that are transcribed from a 10kb region of chromosome 12qF1 are upregulated in the brain and muscle. Interestingly, two other snoRNA: *Gm22962* and *Gm24598* followed the same tissue-specificity pattern, despite being transcribed from other chromosomes (9qC and XqA7.1 respectively). While present at low levels, we also identified *Snora35* (MBI-36) and Snord116 (MBII-85), known to be involved in neurodevelopmental disorders (Cavaille et al., 2000; Gallagher et al., 2002), to be brain exclusive (*TSI*=1). We observed high levels of *Snord17* and *Snord15a* in the spleen and bone marrow, and lower levels in other tissues. These snoRNAs have been previously reported among upregulated genes in bacterial infection of soft tissues (Brady et al., 2015; Podolska et al., 2012), suggesting the association of these transcripts with immune cells. We found several snoRNAs present mainly in the pancreas such as *Snord123* (**Figure S2C**), located 3kb upstream of the pancreatic cancer-associated *Sema5a* gene, and *Gm22888* (**Figure 2B**) located within introns of Ubap2 gene. We also identified a large number of other snoRNA, the exact function of which is still unknown, to be enriched in either one or multiple tissues (**Figure S2C**), among which are *Snord53* and *Gm24339*, *Gm26448*, *Snora73a*, *Snord104* in lymphoid tissues, and *Snord34*, *Gm24837* – in testes. Finally, we show that some snoRNA, such as *Snord70* and *Snord66*, which are often used as normalization controls in qPCR-based assays (Chen et al., 2012; Emde et al., 2015), are also expressed in a tissue-biased manner (**Figure S2C**).

### Tissue-specific expression of *Rny*, *Terc* and other ncRNA

Analyzing the levels of other ncRNA classes, we found that brain contains high levels of *Rny1* compared to other tissues (**Figure 2B**). We also observed that the levels of another transcript from the same class, *Rny3*, are elevated in pancreas, brain and kidney (**Figure S2D)**. The precise biological function of *Rny1* and *Rny3* is so far undefined, although they have been suggested to maintain RNA stability (Hizir et al., 2018; Kowalski and Krude, 2015).

Interestingly, we detected the presence of telomerase RNA component (*Terc*) in analyzed somatic tissues, with highest levels seen in the bone marrow and spleen. Together with previous reports that identify telomerase activity in hematopoietic cells (Morrison et al., 1996) and show *Terc*+ cells to secrete inflammatory cytokines (Liu et al., 2019; Morrison et al., 1996), our data suggests that *Terc* is specific to cells of hematopoietic origin. Among other ncRNA types that we found to be differentially expressed across profiled tissues are snRNA and scaRNA, both known to be involved in the regulation of splicing events (Matera and Wang, 2014). We observed that Rnu11 and Scarna6 are preferentially found in lymphoid tissues, while three snRNA of unverified function, *Gm25793, Gm22448, Gm23814*, are specific to the brain (**Figure S2D**).

### Tissue-specific miRNA

We found ~400 miRNAs differentially expressed across profiled tissues (**Table S3**). Out of these 400, nearly one quarter are specific to the brain, with some being uniquely expressed within the tissue (**Figure S3A**). We identified both well-described brain-specific miRNAs, such as *mir-9*, *mir-124*, *mir-219*, *mir-338* (Landgraf et al., 2007; Ludwig et al., 2016; McCall et al., 2017) as well as those which are missing from existing catalogs, such as *mir-666*, *mir-878*, *mir-433*, etc. (**Figure 2B and Figure S3A**). Examples of other miRNAs previously unknown to be tissue-specific include *mir-499* in the heart, *mir-3073b* in the kidney, *mir-215* and *mir-194* in the intestine (**Figure S3A**). We also observed multiple miRNAs being present in several tissues but absent in others, reflecting the cellular composition of the tissues. Surprisingly, we also found a few miRNAs, such as *mir-134*, *mir-182*, *mir-376c*, *mir299a*, mir-*3061* and *mir-7068* (**Figure S3A)** to be shared solely between muscle, brain and pancreas, which, in turn, do not contain any evident shared cell types that are absent in other tissues. Independently, unsupervised clustering of the top 400 most differentially expressed ncRNA in our dataset also revealed that two out of three identified clusters comprise ncRNA genes which are upregulated in the above-mentioned three tissues (**Figure S3B**). Altogether, these findings suggest an important role of small ncRNA in maintaining a specific function within the brain, pancreas and muscle, which could, for example, be ion transport or exocytosis.

### Tissue-specific arm selection of miRNA

Assessing the overall abundance of 5p or 3p arms of miRNA across tissues, we found no significant bias in strand selection (**Figure S4A**). For many miRNAs we generally observed the dominance of either 3p or 5p arm, however, for some we also detected high levels of both arms present in one or multiple tissues (**Figures 2C-D and Figure S5**). Nonetheless, we found that ~5% of all miRNAs switch their arm preference between tissues. Some of them, like *mir-337*, *mir-106b*, *mir-26b*, are represented by both arms in certain tissues while only by one of the arms in other (**Figure 2D and Figure S5**). More striking examples of complete arm switching from one tissue to another are *mir-141* and *mir-350* (**Figure 2D)**. *miR-141-5p* but not *-3p* is present in the brain and the other way around in the testes, while both arms are found in the pancreas and intestine. In the case of *mir-350,* both arms are detected in the bone marrow, brain and spleen, while only 5p arm is present in the heart, lung, muscle and 3p arm in the pancreas (**Figure 2D**). This highlights the complexity of tissue-dependent miRNA biogenesis and indicates that the phenomena of miRNA arm switching, so far only observed in cancer, extends to healthy mammalian tissues (Chen et al., 2018a; Pinel et al., 2019; Ro et al., 2007).

### Ubiquitous ncRNA transcripts

We detected many ubiquitously expressed ncRNAs across tissues (**Figure S4B**). Among these are ncRNAs known to be expressed in a large number of cell-types, such as *let-7d-3p*, *miR-320-3p* (de Rie et al., 2017), ncRNAs the cell-type specificity of which is still unknown, such as *Snord4a* and *Snord55*, as well as those known to be expressed in the cell types that are abundant in all tissues (like endothelial *miR-151-5p*) (**Figure S4B**).

### Novel miRNAs

We have recently demonstrated that the current miRbase annotation of mammalian miRNA remains incomplete but can be readily expanded with the help of emerging small RNA-seq data (Alles et al., 2019). To search for novel miRNA in our data, we first processed all unmapped reads using miRDeep2 (Friedländer et al., 2012) and selected 473 genomic regions harboring a putative miRNA gene supported by at least ten sequencing reads. To refine this list we employed three parallel strategies: 1) we searched for the presence of the putative miRNA in 141 public Argonaute CLIP-seq (AGO-CLIP) datasets from various mouse cell- and tissue-types, 2) we performed a literature and database search for prior mentions of the putative miRNA, and 3) we ran an RT-qPCR validation of selected candidates. Analysis of AGO-CLIP data showed evidence for 214 out of 473 candidates (total > 5 counts). We also found that 87 out of 473 novel miRNA were previously reported within other studies (Dhahbi et al., 2013a; Fehlmann et al., 2018; Javed et al., 2014; Metpally et al., 2013; Sundaram, 2017) (**Figure 3A**). Importantly, 52 novel miRNAs identified by this and previous studies were not present in the AGO-CLIP data (**Table S4**). The RT-qPCR quantification of two miRNAs selected from this list, *17_11530* and *7_16137*, in turn, confirmed the existence of these transcripts (**Figure S6A**). On the other hand, we identified novel miRNAs (*17_8620*, *4_6440*, *9_15723*) that are supported by AGO-CLIP data, prior reports or both, but for which we could not confirm the existence through RT-qPCR (**Figure S6A**). We also found a novel miRNA that, among the three validation methods, was only verified through RT-qPCR. Interestingly, the genomic coordinates of this miRNA, *14_6588*, matched the coordinates of another, annotated one, *mir-802a*. Unlike *mir-802a*, however, *14_6588* is transcribed from the negative DNA strand and is only present in the brain (**Figure S6B**). Altogether, by comparing the miRNA levels measured through RT-qPCR with the tissue transcript abundance identified by small RNA-seq, we validated 12 novel miRNAs that were either also reported by others (**Figure 3B and Figure S5A**) or uniquely identified in the present study (**Figure 3C-D, Figure S6A**).

**Figure 3.**
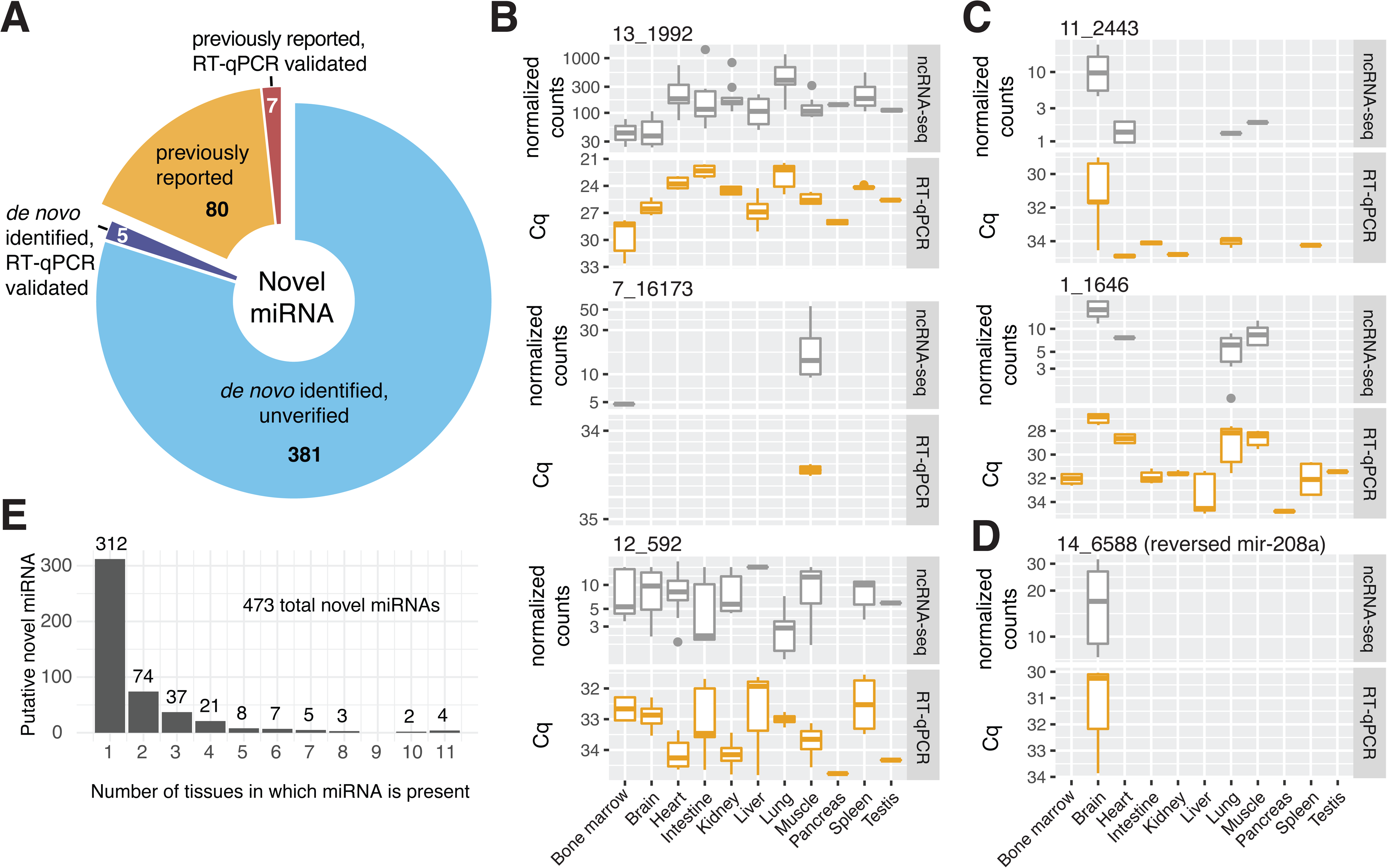
Novel miRNA detected in the present study. **(A) Pie chart** of predicted and verified novel miRNA. **(B) Examples** of RT-qPCR verified novel miRNAs identified by this and previous studies. The levels of novel miRNA were determined from small RNA-seq data (DESeq2 normalized counts) and through RT-qPCR (Cq, adjusted for the sample-to-sample variability using cel-mir-39 spike in control). **(C) Same as C** but for novel miRNAs detected uniquely within the present study. **(D) RT-qPCR** verified novel miRNA, 14_6588, transcribed from the negative strand of mir-208a. **(E) Tissue** specificity of putative novel miRNA.

We found that the majority of putative miRNAs are present in only one tissue (312), but a small number (4) are found in all eleven tissues (**Figure 3E**). Principal component analysis on the newly identified miRNAs, supported by at least 50 reads, showed a clear separation of brain, lung and muscle from other tissues based on the expression values. Similar to annotated transcripts, novel miRNAs demonstrate a spectrum of tissue specificity with some being ubiquitously expressed, while others are only present in one tissue (**Figure S6C**). Differential expression analysis on putative novel miRNAs identified six miRNAs to be also expressed in a sex-specific manner. Strikingly, all six were male-dominant, with one of them even found to be consistently upregulated in two tissues, male muscle and pancreas (**Figure S6D**). We speculate that the prevalence of male-specific novel miRNAs identified in our study reflects the inconsistent sampling of both sexes by prior murine miRNA research.

### Tissue-resident tRNA fragments

About a quarter of our small RNA-seq libraries consisted of tRFs –fragments of either mature of precursor tRNA molecules enzymatically cleaved by angiogenin (Ang), Dicer, RNaseZ and RNaseP (Kumar et al., 2016; Lee et al., 2009; Maute et al., 2013; Telonis et al., 2015a; Thompson and Parker, 2009) (**Figure 4A**). Several recent studies have demonstrated the implication of tRNA-derived ncRNAs in various biological functions, such as cell growth and proliferation (Gebetsberger et al., 2017; Goodarzi et al., 2015; Haussecker et al., 2010; Yamasaki et al., 2009) as well as its uneven distribution across human and primate tissues (Jehn et al., 2019; Telonis et al., 2015a).

**Figure 4.**
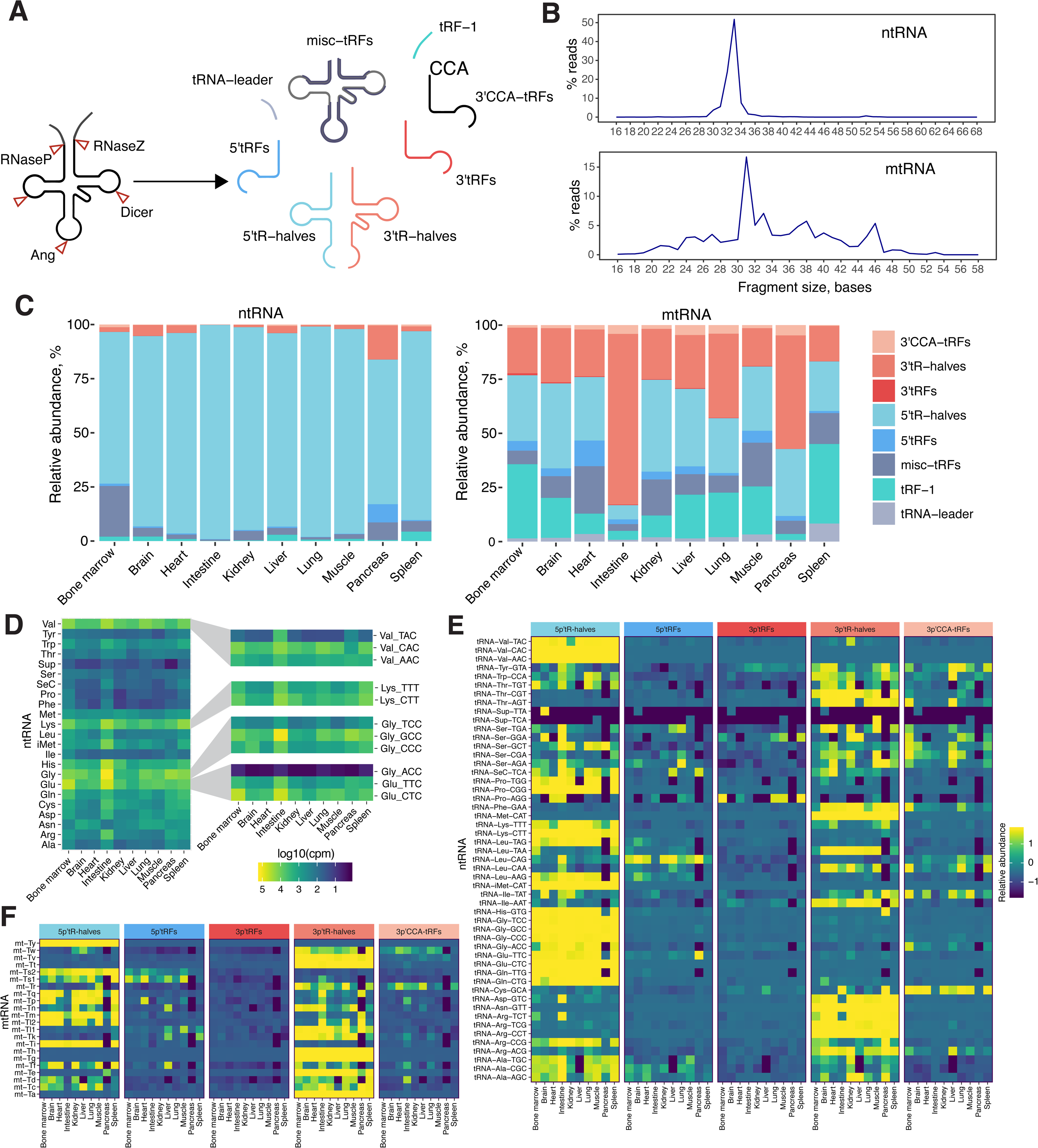
tRFs detected in mouse tissues. **(A) Schematic depiction** of tRNA cleavage and the resulting fragments. **(B) Average tRF length** identified across tissues for either ntRNA-derived (top) or mtRNA-derived (bottom) fragments. **(C) Fragment type abundance** across tissues. **(D) Heatmap** of tRF levels in tissues. For each of the 23 tRNA types the sum among its tRFs is plotted. **(E) Heatmap** of relative abundance of ntRNA fragment types (5’-tR-halves, 3’-tR-halves, 5’-tRFs, 3’ tRFs or 3’-CCA-tRFs) for each tRNA isoacceptor across ten somatic tissues. Relative abundance is represented by row-wise scaled fractionated scores of tRFs computed by *unitas*. **(F)** Same as E but for mtRNA.

‘Exact tRNA multi-mapping’ (Telonis et al., 2015a) of these fragments to the mouse genome revealed the presence of tRFs of various sizes. Interestingly, consistent with a previous report on tRFs in human cell lines (Telonis et al., 2015a), we observed a major difference in the size of fragments originating from either nuclearly- or mitochondrially encoded tRNA (ntRNA and mtRNA respectively). While the majority of ntRNA fragments were 33 nt long, mtRNA fragments spanned a large size range of 18 to 54 nt (**Figure 4B and Figure S7A**). This distinct pattern of fragment sizes reflected the bias in the amounts of tRF types originating from nt- and mtRNA (**Figure 4C and Figure S7B**). We observed that the distribution of nuclear tRFs was largely skewed towards 5’tR-halves, generated by the cleavage in the anticodon loops of mature tRNA. However, within mitochondrial tRFs we identified a more uniform representation of cleaved fragments. Furthermore, we found that the relative abundance of tRF types, within both ntRNA and mtRNA space, is not constant, but varies across tissues (**Figure 4C-D**). The tissue-type differences are also present across different tRNA isoacceptors and evens its anticodons (**Figure 4D and Figure S7B**). In the case of nuclear tRFs the vast majority of fragments in each tissue was attributed to glycine, glutamine, valine and lysine tRNA, with the intestine containing the largest amounts of the respective 5’tR-halves. Since the abundance of these specific fragments has been shown to correlate with the levels of functional angiogenin in the cell (Thomas et al., 2018), we speculate that the biological explanation of the intestine yielding high levels of tRFs is due to the activity of Ang4, one of the five Ang proteins in mouse, highly expressed in Paneth and Goblet cells of the intestinal epithelium (Forman et al., 2012; Hooper et al., 2003). We cannot, however, rule out the possibility that some of these fragments were obtained from a mouse diet enriched with nutrients of mammalian origin.

For many ntRFs the distribution between 3’- and 5’-, tRF and tR-halves was surprisingly shifted towards one form, i.e. one fragment type was present at higher amounts than others (**Figure 4E-F**). For the majority of fragments, we found 5’tR- or 3’tR-halves to be the most dominant fragment type. However, in the rare cases, we found 5’- or 3’tRFs to dominate other fragments in a tissue-specific manner. An example of such a fragment is 5’tRF Glu-TTC, which we found to be enriched in the pancreas, compared to other tissues that mostly contained Glu-TTC 5’tR-halves. mtRFs followed a similar trend of fragment shift. We found 5’tRFs of proline-transferring mt-Tp in the heart and 5’tRFs of asparagine-transferring mt-Tn in the liver, while within other tissues we detected different fragment types of these tRNAs (**Figure 4F**).

### miRNAs are expressed in a sex-specific manner

Several groups have observed a sex bias in the levels of miRNA in blood, cancer tissues and human lymphoblastoid cell lines (Guo et al., 2017; Kolhe et al., 2017; Loher et al., 2014; Meder et al., 2014). To investigate whether this phenomenon extends to healthy tissues, we compared the ncRNA levels within each tissue coming from either female or male mice. Among ~6000 genes assigned to various ncRNA classes, we identified several miRNAs to be differentially expressed between females and males (at FRD<0.01) (**Figure 5A**). Some of them are globally sexually-dimorphic, while the majority are sex-biased only within a specific tissue. In each somatic tissue, except pancreas, we identified at least two differentially expressed miRNAs between sexes (log2FoldChange > 1, normalized counts > 100, FDR < 0.01) (**Figure S8A-B**). Kidney and lung contained the highest number of sex-biased miRNAs (27 and 18 respectively), while only two were detected in the heart, five in the muscle and seven in the brain **(Figure 5B and Figure S8A).** Three out of eight female-dominant miRNAs: *mir-182*, *mir-148a* and *mir-145a*, were also shown previously to be estrogen regulated (Klinge, 2009) while another miRNA, *mir-340*, was reported to be downregulated in response to elevated androgen levels (Fletcher et al., 2014). Interestingly, we also found that four out of five male-specific miRNAs in the brain are transcribed from a 5kb region of imprinted Dlk1-Dio3 locus on chromosome 12 (Glazov et al., 2008) (**Figure 5C**).

**Figure 5.**
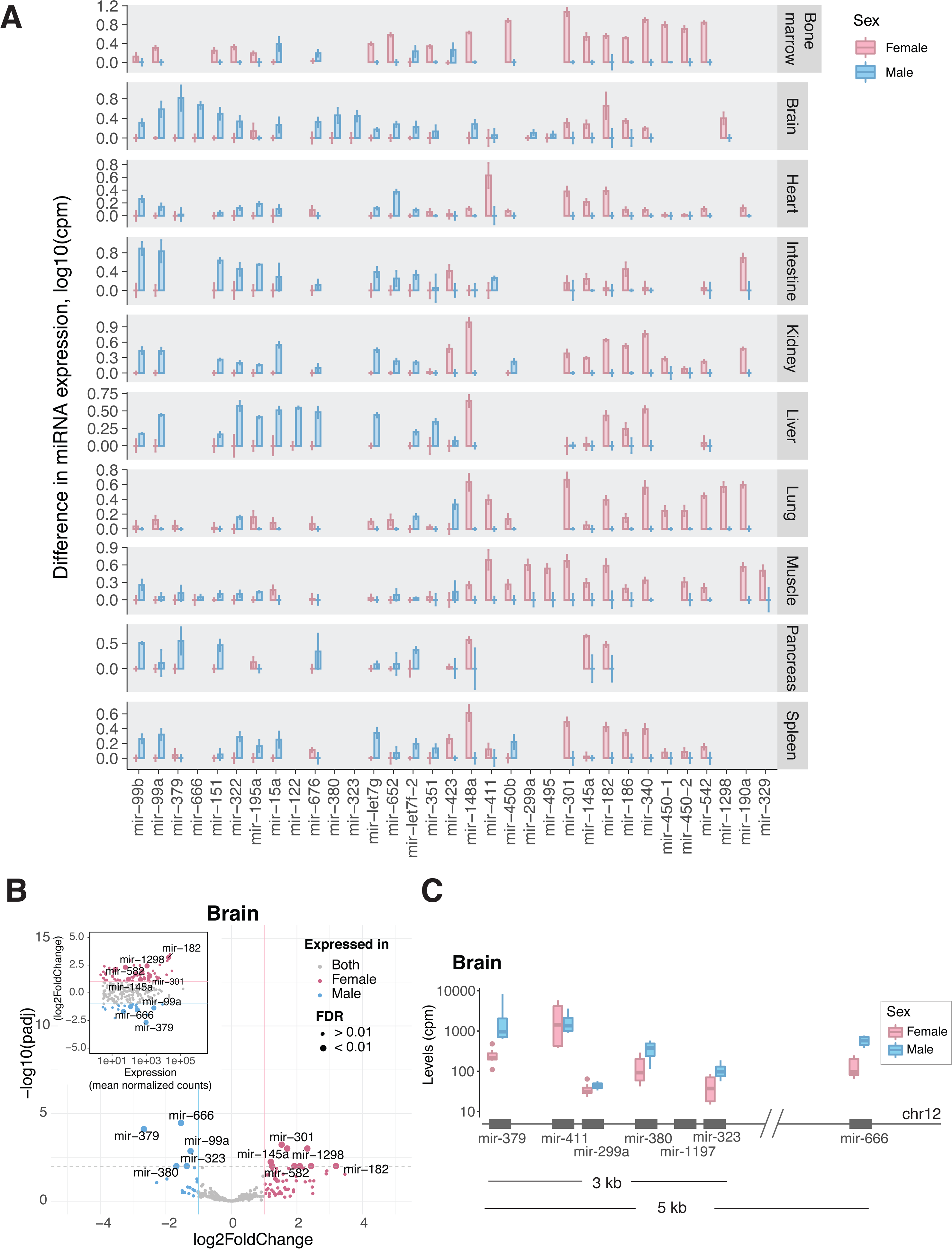
Sex-specific miRNA expression. **(A) Sex-dimorphic miRNAs** identified in this study. Y-axis represents the log10 of the difference between mean miRNA levels computed for males and females in each tissues (in cpm). Error bars denote standard deviation of miRNA levels arcoss a tissue within each sex. **(B) Volcano plot** showing miRNAs differentially expressed between the female and the male brain. **(C) Expression** and gemonic location of male-biased miRNA in the brain.

Given the innate ability of miRNA to lower the levels of target mRNA (Guo et al., 2010), we hypothesized that the levels of protein-coding transcripts targeted by sex-biased miRNA would also differ across male and female tissues. To test this hypothesis, we correlated the expression of sex-biased miRNAs with the levels of their respective target mRNAs across profiled tissues (*Methods*). Among the anticorrelated targets (*r_s_* < −0.8, FDR < 0.1) we identified two genes previously shown to be sexually dimorphic (**Figure S8C**). Specifically, we found *miR-423*, upregulated in male lung and bone marrow, to negatively correlate with its target – estrogen-related receptor gamma (*Esrrg*) (*r_s_*= −0.9, FDR < 0.1), and female-specific *miR-340* to negatively correlate with androgen-associated ectodysplasin A2 receptor (*Eda2r*) (Prodi et al., 2008).

### ncRNA-based tissue classification

It is natural to wonder whether the observed variation in ncRNA expression across tissues (**Figure 2B-C**) would be sufficient to accurately predict the tissue type based solely on small RNA-seq data. To address this question, we set out to construct an algorithm that can learn the characteristics of a healthy tissue from the data reported in the current study and make predictions on new data sets. We limited our analysis to miRNA, since high-throughput tissue data for other ncRNA types is not available. We first trained a support vector machine (SVM) model (Cortes and Vapnik, 1995) on data sets generated in this study, each containing the expression scores for 1973 miRNAs (**Figure S9A**). As a validation dataset we used available miRNA-seq data released by the ENCODE consortium for multiple mouse tissues (Dunham et al., 2012). Notably, the ENCODE datasets contained data generated for the postnatal and embryonic life stages, as opposed to the adult stage profiled in the current study (**Table S5**). Nonetheless, our SVM model accurately classified postnatal forebrain, midbrain, hindbrain and neural tube as brain tissue, as well as accurately inferred the tissue types for heart, intestine, kidney, liver, muscle samples, yielding an overall accuracy of 0.96 (*Methods*). For the embryonic tissues, however, our model was able to only reach an accuracy of 0.69. This was mainly due to inability of the model to correctly classify liver tissues and instead assigning them to bone marrow (**Figure 6A**). Strikingly, in this case our model accurately predicted the hematopoietic composition of the organ, known to shift from the liver at the embryonic stages to the bone marrow in adulthood (Baron et al., 2012), rather than the tissue type itself. Furthermore, we identified hematopoiesis-associated *miR-150* and *miR-155* (Bissels et al., 2012) to have highest weights among the features defining the bone marrow in our model (**Figure S9B**).

**Figure 6.**
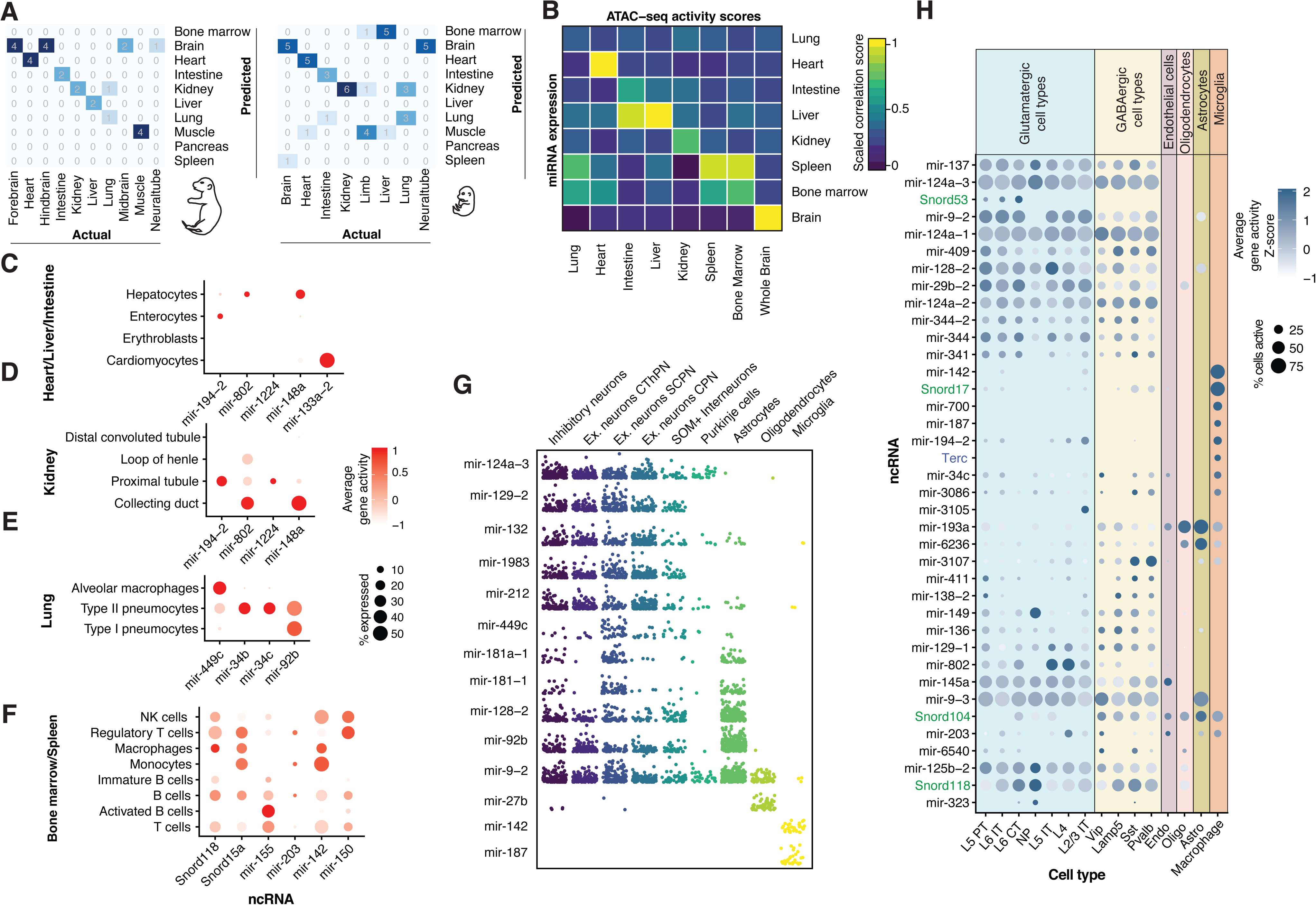
**(A) Confusion matrices** obtained from SVM tissue classifier for postnatal and embryonic datasets. **(B) Correlation** between small RNA-seq scores derived for top 400 tissue-specific ncRNA genes with the respective activity scores obtained from Mouse ATAC atlas. **(C-F) Average gene activity** scores of tissue-specific ncRNAs within each cell type resident to the respective tissue. Gene activity scores for the ncRNAs of interest were retrieved from the Mouse ATAC atlas. **(G) Log10 gene activity** scores of brain-specific miRNA across individual cells. **(H) Gene activity** scores computed for brain-specific ncRNA from the droplet-based scATAC-seq data (10XGenomics, see *Methods*).

We next asked how the identified tissue expression patterns compare to those of individual cell types. To investigate that, we correlated our data to the miRNA data generated for primary mammalian cells by FANTOM5 consortium (de Rie et al., 2017). Comparing mouse samples first, we found that FANTOM5 embryonic and neonatal cerebellum tissues strongly correlated with our brain samples (*r_s_*=0.89-0.9), while erythroid cells had the strongest correlation with spleen and bone marrow (*r_s_*=0.93) (**Figure S9C**). To perform a comparison with human samples, we focused on the expression scores of 531 orthologs detected in both the current study and the FANTOM5 samples (**Figure S9D**). Spearman correlation coefficients reflected the cell-type composition of tissues (**Figure S9E**). As such, we observed that mouse bone marrow and spleen had the highest correlation with human B-cells, T-cells, dendritic cells and macrophages (0.5<*r_s_*<0.6), muscle correlated the most with myoblasts and myotubes (*r_s_*=0.47), while brain – with neural stem cells, spinal cord, pineal and pituitary glands (*r_s_*=0.49) **(Figure S9E)**.

### Integration of small RNA-seq and scATAC-seq data

Finally, to deconvolute the complex non-coding tissue profiles and identify the cell types that contribute the observed tissue-specific ncRNAs, we integrated the sequencing data generated within our study with a previously published single-cell ATAC-seq atlas – a catalog of single-cell chromatin accessibility profiles across various cell types (Cusanovich et al., 2018). First, we compared the expression scores predicted through ATAC-seq measurements with our estimates of ncRNA expression, derived from small RNA-seq. Using top 400 ncRNAs identified in our analysis as tissue-specific, we correlated average ATAC-seq activity scores and ncRNA levels across 8 tissues for which both types of data were available. We observed a strong correlation of both measurements for the brain, liver and heart, while weaker correlation for kidney and mixed scores between spleen, bone marrow and lung (**Figure 6B**). We found, however, that within each tissue we could attribute the cell-type of origin to a number of identified tissue-specific ncRNA. For example, in agreement with previous studies, we could identify that muscle-specific *mir-133a-2* is expressed in cardiomyocytes, while *mir-148a* is expressed in hepatocytes and duct cells in liver and kidney (**Figure 6C-D**) and *mir-194-2* comes from the enterocytes in the gut (McCall et al., 2017; Peck et al., 2017; de Rie et al., 2017). In addition, we found that in the lung, *mir-449c* in expressed in alveolar macrophages, *mir-34b* and *mir-34c* in type II pneumocytes (**Figure 6E**), *mir-155* is found in B cells and even correlates with its maturation status (**Figure 6F**). Among brain-specific ncRNA, we identified *mir-187* and *mir-142* to be expressed in microglia, *mir-27b* – in oligodendrocytes, *mir-124a-3*, *mir-1983*, *mir-212* and others – in neurons (**Figures 6G**). For the majority of brain-specific ncRNA identified in our study, however, due to resolution limitations of the Mouse ATAC atlas data, we were unable to unambiguously conclude the cell-type of origin. To overcome that, we analyzed a complimentary single-cell ATAC-seq dataset, generated specifically from the mouse adult brain (obtained from 10X Genomics, see ***Methods***) and mapped the activity of the brain-specific ncRNA to 15 cell-types in the adult mouse brain annotated by the Allen Brain Atlas (Lein et al., 2007). This analysis revealed that among the brain-specific ncRNA, many are potentially expressed solely in neurons, with some even being predominantly present in either glutamatergic (*Snord53*, *mir-802*) or GABAergic (*mir-3107*) neurons (**Figure 6H**). Among glia-specific ncRNA, we identified *Snord17* and *mir-700* in macrophages as well as *mir-193a* and *mir-6236* – in astrocytes and oligodendrocytes.

## DISCUSSION

Small non-coding RNA plays an indispensable role in shaping cellular identity in health and disease by orchestrating vital cellular processes and altering the expression of protein-coding genes (Kuscu et al., 2018; Matera et al., 2007; Qureshi and Mehler, 2012). Recent efforts in profiling of the most studied types of small ncRNA, miRNA, across cells and tissues demonstrated the existence of tissue- and cell type-specific short non-coding transcripts. (Landgraf et al., 2007; Londin et al., 2015a; Ludwig et al., 2016; McCall et al., 2017; de Rie et al., 2017). In this work, we show that this phenomenon extends beyond one ncRNA class and involves not only tissue-specific but also sex-specific ncRNA expression. The present resource demonstrates that each healthy mammalian tissue carries a unique non-coding signature, contributed by well-understood RNA types as well as by the less studied ones.

By analyzing the expression of several classes of ncRNA we discovered that nearly ~900 various transcripts contribute to the unique non-coding tissue profile. Moreover, we identified that in addition to variable transcription levels and posttranscriptional modifications (Ebhardt et al., 2009), non-coding tissue-specificity is achieved through an unknown mechanism of selective RNA retention. While at this point we are unable to judge the functional significance of this phenomenon, we discovered that, even between healthy tissues, certain miRNA undergo so-called “arm switching” – a process previously thought to be strictly pathogenic in mammals (Chen et al., 2018b; Lin et al., 2018; Wilson et al., 2015). Among other ncRNA class, tRFs, we observed a selective enrichment of certain fragment types over other, happening in both, gene- and tissue-specific manner. Taken together with previous observations (Anderson and Ivanov, 2014; Cole et al., 2009; Dhahbi et al., 2013b; Sharma et al., 2016; Telonis et al., 2015b), this finding raises additional questions regarding the biogenesis pathways of tRFs as well as their tissue-specific function.

Within our study we also report new tissue-specific miRNAs not identified in previous studies (**Figures 3 and Figure S4, Table S6**). The validation process of these novel miRNAs brought to light several important observations. First, we noted that the AGO-CLIP, while often used as a “gold standard” of miRNA validation (Londin et al., 2015b; McCall et al., 2017), in fact, does not support the existence of many miRNA independently detected within RNA-seq datasets or directly validated through RT-qPCR. The gap between AGO-CLIP and small RNA-seq data in terms of data quality, diversity and depth suggests that validating against AGO-CLIP data may not be the optimal approach for miRNA discovery. Instead, one could search for the evidence of miRNA expression within publicly available RNA-seq data as a first thresholding step (Backes et al., 2016). Second, it is important to consider that the genomic location of a novel miRNA might match with that of a previously annotated one, while the molecule itself could be transcribed from an opposite strand. We observed this phenomenon on the *de novo* identified miRNA 14_6588, whose coordinates strictly overlap with *mir-802a* and that is only present in the brain.

More than 60% of protein-coding genes in mammalian genome harbor predicted miRNA target sites (Friedman et al., 2008). However, only a handful of them are *bona fide* miRNA targets, while up to 70% are falsely assigned by prediction algorithms (Agarwal et al., 2015; Betel et al., 2010; Dweep and Gretz, 2015). Currently, the validation of miRNA:mRNA interactions is still mainly based on low-throughput labor-intensive approaches such as knock-down or over-expression assays (Thompson et al., 2015) and thus has been only done for a limited number of miRNAs. Growing amounts of both, mRNA- and miRNA-seq data, generated for various cell and tissue types, now provide us with a possibility to narrow down the list of putative targets by identifying those that go down with elevated miRNA levels (Guo et al., 2010). Here, using our tissue miRNA-seq dataset and publicly available mRNA-seq data, we demonstrate the implementation of this approach. By correlating the expression of miRNA:target pairs across tissues, we show that at least half of the predicted targets are not affected by increasing miRNA levels. In parallel, we were able identify targets that show a strong negative correlation with miRNAs. On the example of sexually dimorphic miRNA we further demonstrate that the levels of some protein-coding transcripts indeed decrease with increased miRNA levels (**Figure S8C**).

miRNA has been previously used to train classifiers capable to differentiate cancer/tissue types (Rasnic et al., 2017; Sherafatian, 2018). Our work demonstrates that machine learning algorithms applied to quantitative miRNA expression estimates also detect changes related to the cell type composition of tissues, such as the shift in hematopoietic cell abundance in the postnatal compared to the fetal liver. Given the emerging evidence of ncRNA stability in the blood and its rapid propagation throughout the body within extracellular vesicles (Bhome et al., 2018), we anticipate that the current space of markers used to non-invasively monitor development (Ngo et al., 2018) could be further expanded to small ncRNA.

Small non-coding RNAs have been long known to regulate the development and function of the brain (Qureshi and Mehler, 2012). Despite the tremendous progress of neuroscience in understanding the regulation of coding genes, surprisingly little is known about cell-type specific small ncRNA in the brain. Even within the available tissue-level ncRNA resources, brain remains one of the most underrepresented tissue. We believe this is mostly due to technical limitations of small RNA sequencing, which is yet to be applicable to single neurons and, so far, still relies on the robust enrichment of certain cell types. Our study finds that brain, in fact, contains the largest number of unique mammalian ncRNA transcripts that are absent in other tissues. However, our knowledge of cell-specific ncRNA expression is not complete and thus for the majority of these identified RNA we could not call the cell type of origin based in the data generated within previous studies offering cell-type resolution (McCall et al., 2017; de Rie et al., 2017). Taking an alternative route and integrating our tissue-level ncRNA measurement with single-cell chromatin accessibility profiles turned out to be surprisingly informative and allowed us to infer the activity of ncRNA within individual neuronal and glial types. While the validation of these cell-specific transcripts through a direct measurement remains highly desirable, the provided ncRNA estimates indicate that ncRNA is another contributor of complexity in the architecture of nervous system.

We found that lung contains the largest number of distinct small ncRNA among eleven profiled tissues. However, in the case of lung, open chromatin data did not provide sufficient resolution for us to infer the cell-types of origin for the majority of the transcripts. This inability to fully explain the roots of tissue complexity points to the need for further characterization of the ncRNAs content of specific cell types or even, similarly to mRNA, that of single cells (Faridani et al., 2016; Trapnell, 2015). This atlas, meanwhile, will hopefully stimulate future small ncRNA studies and serve as a powerful resource of ncRNA tissue identity for fundamental and clinical research.

## Acknowledgements

We thank Dylan Henderson for assistance in RNA extraction and library preparation. Norma Neff and Jennifer Okamoto for sequencing expertise. Geoff Stanley and Kiran Kocherlakota for kind advice in tissue dissection and preservation. Jennifer Okamoto and Norma Neff for the assistance in sequencing of the small RNA-seq libraries. This study was supported by Howard Hughes Medical Institute and Chan Zuckerberg Biohub. A.I. was supported by the Swiss National Foundation Early PostDoc Mobility Fellowship.

## Author Contributions

A.I. designed and performed the experiments and data analysis. T.F. and A.K. supported the data analysis. A.I. and S.Q. interpreted the data and wrote the manuscript.

## METHODS

### Subject details

#### Animals

All procedures followed animal care and biosafety guidelines approved by Stanford University’s Administrative Panel on Laboratory Animal Care and Administrative Panel of Biosafety. Wild type C57BL/6J mice, 4 males and 10 females, aged ~3 month old were used (**Table S1**).

#### Tissue handling and RNA extraction

Upon collection, tissue samples were submerged and preserved at −80C in RNAlater stabilization solution (ThermoFisher cat # AM7021) until further processing. Total RNA was isolated from ~ 100 mg of tissue using Qiagen miRNeasy mini kit (cat # 217004) and the Qiagen tissue lyser using 5 mm stainless steel beads. RNA integrity was assesses using Agilent Bioanalyzer using RNA 6000 pico kit (Agilent Technologies cat # 5067-1513).

#### Library preparation and sequencing

Short RNA libraries were prepared following the Illumina TruSeq Small RNA Library Preparation kit (cat # RS-200-0012, RS-200-0024, RS-200-0036, RS-200-0048) according to the manufacturer’s protocol and size-selected using Pippin Prep 3% Agarose Gel Cassette (Safe Science) in a range 135 bp – 250 bp. Samples were pooled in batches of 48 and sequenced using the Illumina NextSeq500 instrument in a single-read, 50 or 75-base mode.

#### Data processing

Sequencing reads were demultiplexed by BaseSpace (Illumina). Reads were trimmed from the adaptor sequences and aligned to the mouse genome (GRCm38) following ENCODE small RNA-seq pipeline (ENCODE Project Consortium, 2012), with minor modifications. We used STAR v2.5.1 (Dobin et al., 2013) with the following parameters -- outFilterMismatchNoverLmax 0.04 --outFilterMatchNmin 16 --outFilterMatchNminOverLread 0 --outFilterScoreMinOverLread 0 --alignIntronMax 1 --outMultimapperOrder Random --clip3pAdapterSeq TGGAATTCTC --clip3pAdapterMMp 0.1. We allowed incremental mismatch: no mismatches in the reads <=25 bases, 1 mismatch in 26-50 bases, 2 in 51-75 bases. Spliced alignment was disabled. We additionally filtered out reads “soft-clipped” at the 5’-end but kept 3’-clipped ones to account for miRNA isoforms and tRNA modifications. We used GENCODE M20 (Frankish et al., 2019) and miRBase v22 (Kozomara and Griffiths-Jones, 2014) annotations to count the number of ncRNA transcripts. For snoRNAs, snRNAs, scaRNA, miscRNA or miRNA quantification, reads were assigned to the respective genes using *featureCounts v 1.6.1* (Liao et al., 2014) with the following parameters -a Mus_musculus.M20.gtf -M –primary -s 1. To account for the reads from both intronic and exonic regions of the protein coding and lincRNAs for each read mapping within two overlapping feature we assigned a count of 1. To account for the multimappers, we used -M -primary option which only counts a “primary” alignment reported by STAR (either to a location with the best mapping score or, in the case of equal multimapping score – to the genomic location randomly chosen as “primary”). This quantification approach largely agreed with the results obtained through mapping and quantification against a short nucleotide library (Lu et al., 2018) (**Figure S10**). However, it proved to be more inclusive for the reads uniquely mapping within the miRNA exon but missing one base at the 5’prime end of the molecule and more strict in counting reads mapping elsewhere in the genome, for which the levels were consistently overestimated by the other method. All reads mapping miRNA arms and the stem loop were used to quantify miRNA expression at the gene level. For tRF quantification, for each library we first extracted reads mapped by STAR to GENCODE-annotated tRNA within the mouse genome (Chan and Lowe, 2016; Gebert et al., 2017). We then ran *unitas (Gebert et al., 2017)* on these reads and used fractionated scores to compute the differences in tRF abundance across tissues.

### Unsupervised clustering and dimensionality reduction analysis

Raw counts were normalized and log-transformed using *DESeq2* package. Batch effects were corrected using *limma* R package (Ritchie et al., 2015). Hierarchical clustering was performed using log_2_ transformed expression values and using complete linkage as distance measure between clusters. We computed Euclidian distances between samples and used these values to perform the *t*-distributed stochastic neighborhood embedding (*t*-SNE) (Laurens Van Der Maaten and Geoffrey Hinton, 2008) with the following parameters: perplexity = 20, initial dimensions of 50 and maximum iteration of 1,000. Transcripts detected in one or more samples with overall log_2_ expression scores <1 were excluded from this analysis.

### Tissue specificity index

To compute the tissue specificity index we used the formula described previously in (Ludwig et al., 2016):

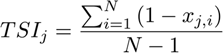

Where *N* is the total number of tissues measured and *x_j,_*_i_ is the expression score of tissue *i* normalized by the maximal expression of any tissue for miRNA *j*.

### Comparison with available miRNA data

To compute Spearman correlation coeffitients between samples generated in the current study and mouse miRNA data generated by FANTOM5 consortium (de Rie et al., 2017) we used *DESeq2*-normalized scores of 2207 annotated miRNAs. To compare the miRNA expression between mouse tissues and human cell types we generated a curated list of orthologous miRNAs that contained maximum two mismatches per ortholog mature miRNA. 531 miRNA passed this criteria and were used to compute Spearman correlation coefficients shown in **Figure S9**.

### Differential expression analysis with DESeq2

We used a likelihood-ratio test (LRT) implemented in *DESeq2* (Love et al., 2014) to compute the significance of each gene in tissue-specific expression. Gene clusters in **Figure S3B** were computed on 250 differentially expressed genes (padj<1E-90 and basemean>3) using *DEGreport* R package (Lorena Pantano). miRNAs differentially expressed between female and male tissues were computed based on uniquely mapping counts (excluding multimappers), using Wald test within *DESeq2*. To test for the NULL hypothesis, we performed a permutation test in which we randomly re-assigned the sex labels to 14 samples across each tissue and plotted the distribution of *DESeq2* p-values computed for the two groups (i.e. female and male) (**Figure S8A**). We used Benjamini-Hochberg-corrected p-values to assess the statistical significance of the computed DE scores (**Figure 5 and S8A**). The differentially expressed miRNAs were visualized on volcano plots, where male- and female-specific miRNAs (adjusted *P*-value < 0.01 and absolute fold change > 1) were labeled accordingly.

### Analysis of correlation between miRNA expression and the expression of its targets

Putative miRNA target genes were extracted from TargetScan, DIANA, miRanda, or mirDB databases (Agarwal et al., 2015; Griffiths-Jones et al., 2007; Paraskevopoulou et al., 2013; Wong and Wang, 2015). Only targetes present in two or more databases were used. The gene expression scores of the respective targets in various tissues were extracted from the ENCODE database (Pennisi, 2012) (**Table S6**). Spearman correlation coefficients were computed between FPKM retrieved from the ENCODE mRNA expression tables and *DESeq2*-normalized miRNA counts across ten profiled tissues using corr.test() function from ‘psych’ R package (Revelle, W.) and threshholded above Benjamini-Hochberg adjusted *P*-value of 0.1 and Spearman correlation coefficient (−0.8< *r_s_*<0.8).

### Identification of candidate novel miRNA

Candidate novel miRNA were identified using miRDeep2 software (Friedländer et al., 2012). Only miRNAs supported by > 5 reads were reported in this study. AGO-CLIP data was mapped to the mouse genome using STAR (same as for small RNA-seq libraries described in the Data Processing section) and the reads falling within the novel miRNA coordinates were counted using *featureCounts*. We counted a novel miRNA as supported if it had >5 AGO-CLIP counts.

To search for the previous mentions of novel miRNAs we looked up their sequences in miRCarta (Backes et al., 2018) and used the Google search engine to query the literature. Candidate miRNAs were ranked by novoMiRank scores, which we computed as described in (Backes et al., 2018). Briefly, for 24 features, comprising sequence, structure and genomic features, z-scores were computed for the high confidence set of mouse miRNA from miRBase v22. To limit large influences of single features, we restricted the absolute z-score values to 3. Next, we determined for every feature the distance of the predicted miRNA to the distribution of the high confidence miRNAs and reported the mean z-score.

For independent validation we preformed RT-qPCR using custom Small RNA Taqman probes (Life Techomologues, cat# 4398987) designed based on the star consensus sequence reported by miRDeep2. We used 0.5 ng of total RNA per tissue sample supplied with Cel-mir-39 spike-in (Qiagen, cat# 339390) to perform the reactions in a final volume of 20uL.

We analyzed tissue- and sex-specificity of novel miRNAs based on transcripts supported by at least 50 sequencing reads across all samples. Statistical analysis and data visualization were performed as described above for annotated miRNAs.

### miRNA-based classifier

We trained the radial kernel SVM model on 136 samples corresponding to different tissue types (**Figure S9A**) using *e1070* (Meyer et al., 2017) R package. We used z-scores of *DESeq2* normalized counts obtained in this study as a train dataset and those obtained from ENCODE miRNA-seq data as test dataset (**Table S4**). We normalized and scaled train and test datasets separately.

To measure the predictive power of each model we used the accuracy measure, calculated as the following:

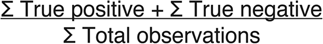

We tuned the SVM model to derive optimal cost and gamma using tune.svm() function and searching within gamma ∈ [2^(−10): 2^10] and cost ∈ [10^(−5):10^3]. We tuned RF model using first random and then grid search, with an evaluation metric set to “Accuracy”. The accuracy was computed using 10-fold cross-validation procedure (Japkowicz and Shah, 2011). The reported accuracy is computed as a mean over the 10 testing sets in which 9-folds are used for training and the held-out fold used as a test set. The R script used to train the models and compute the predictions is included in the supplement.

### Comparison with scATAC-seq data

To compute and plot the correlations of small RNA-seq with scATAC-seq (**Figure 6B-G**) we loaded chromatin accessibility scores reported in Cusanovich et al., 2018, in *Seurat v3,* normalized, scaled and averaged per cell- or tissue-type. To compute the accessibility scores for the brain specific ncRNA in **Figure 6H** have used scATAC-seq data generated by 10XGenomics for the mouse adult brain (https://www.10xgenomics.com/10x-university/single-cell-atac/) with the Chromium Single Cell ATAC platform, and demultiplexed and pre-processed with the single-cell ATAC Cell Ranger platform. Using *Seurat v3* we clustered the cells and merged them with Allen Brain Atlas single-cell RNA-seq data (Lein et al., 2007) for the further transfer of cell annotation labels. We computed the activity scores for brain-specific ncRNA identified through small RNA-seq using *cicero* (Pliner et al., 2018).

### Data availability

The datasets generated and analyzed in the study are available in the NCBI Gene Expression Omnibus (GEO) under the entry GEO:GSE119661.

## Supplementary figures

**Figure S1.**
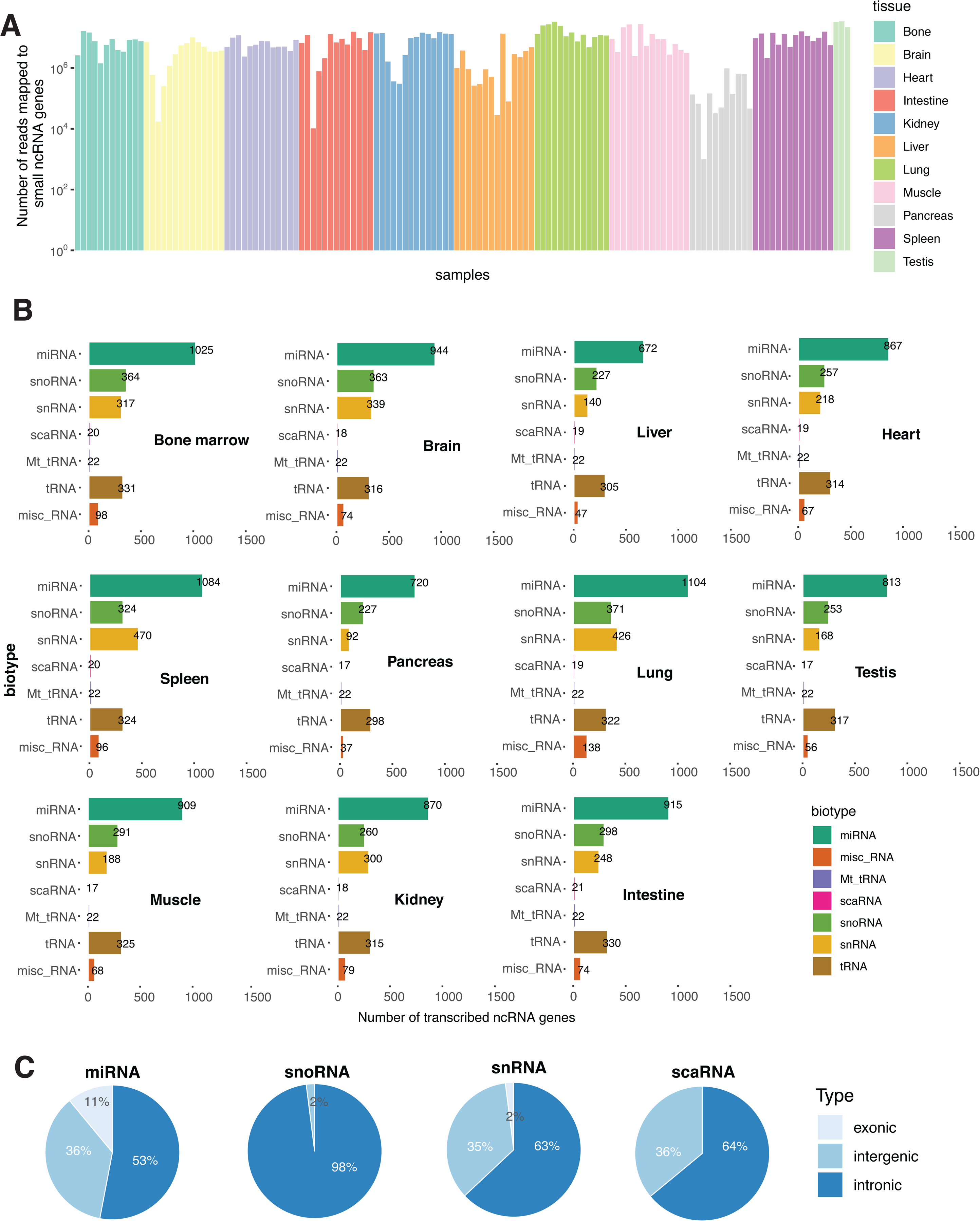
Read statistics. (A) Number of reads mapping to small ncRNA genes across samples. (B) Number of unique ncRNAs detected in each tissue. Numbers indicate detected miRNA, snoRNA, snRNA, scaRNA, Mt_tRNA annotated in GENCODE M20, as well as high-confidence tRNA listed in GtRNAdb. (C) Genomic localization of identified ncRNAs with respect to protein-coding genes.

**Figure S2.**
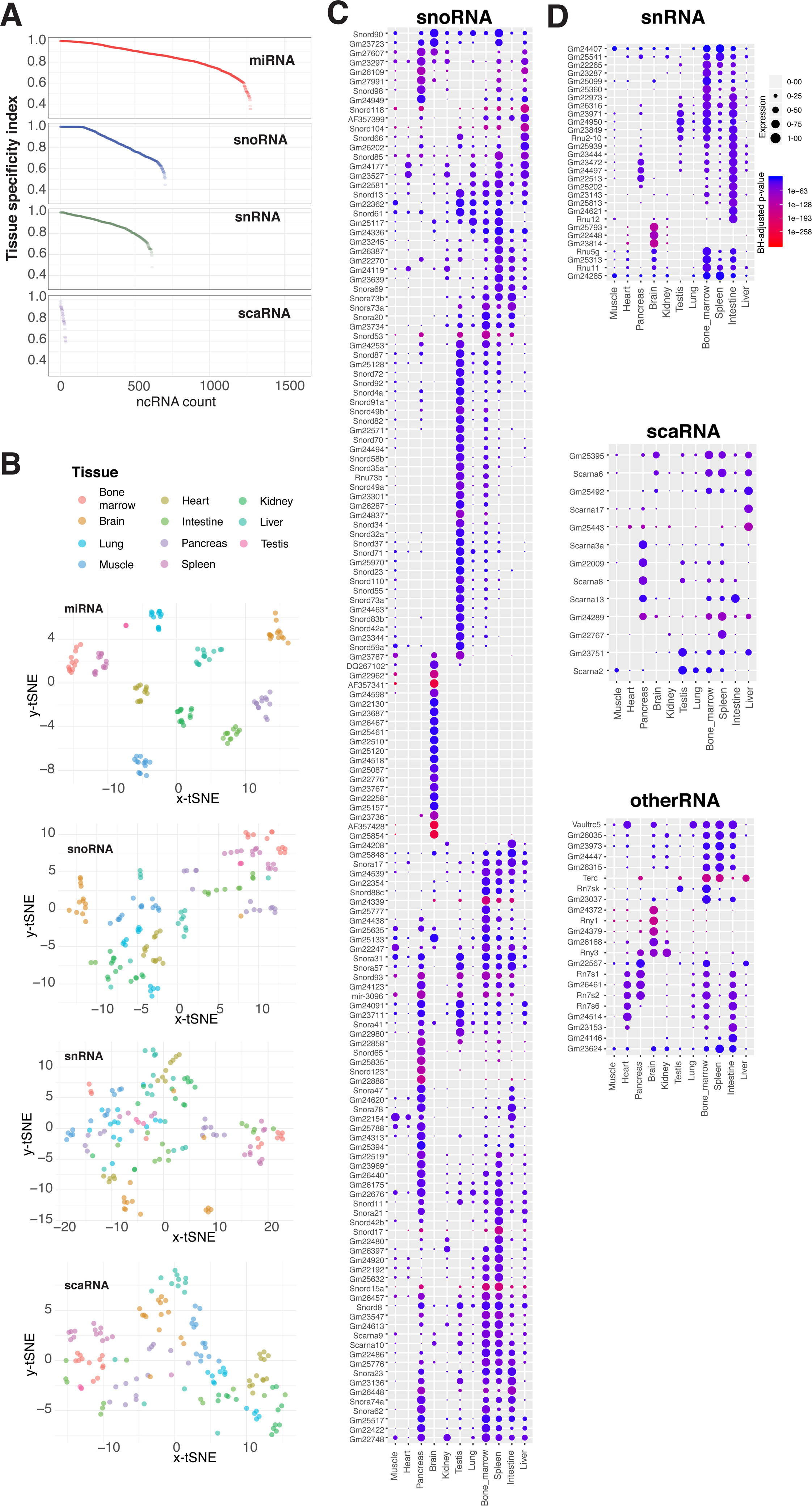
Tissue-specific ncRNA. (A) TSI of miRNAs, snoRNAs, snRNAs and scaRNAs. (B) t-SNE projection of expression values for each ncRNAs class. (C) Expression of top 150 tissue-specific GENCODE-annotated snoRNA (BH-adjusted <0.01 in LRT test) across tissues. (D) Same as (C) for snRNA, scaRNA and other small ncRNA.

**Figure S3.**
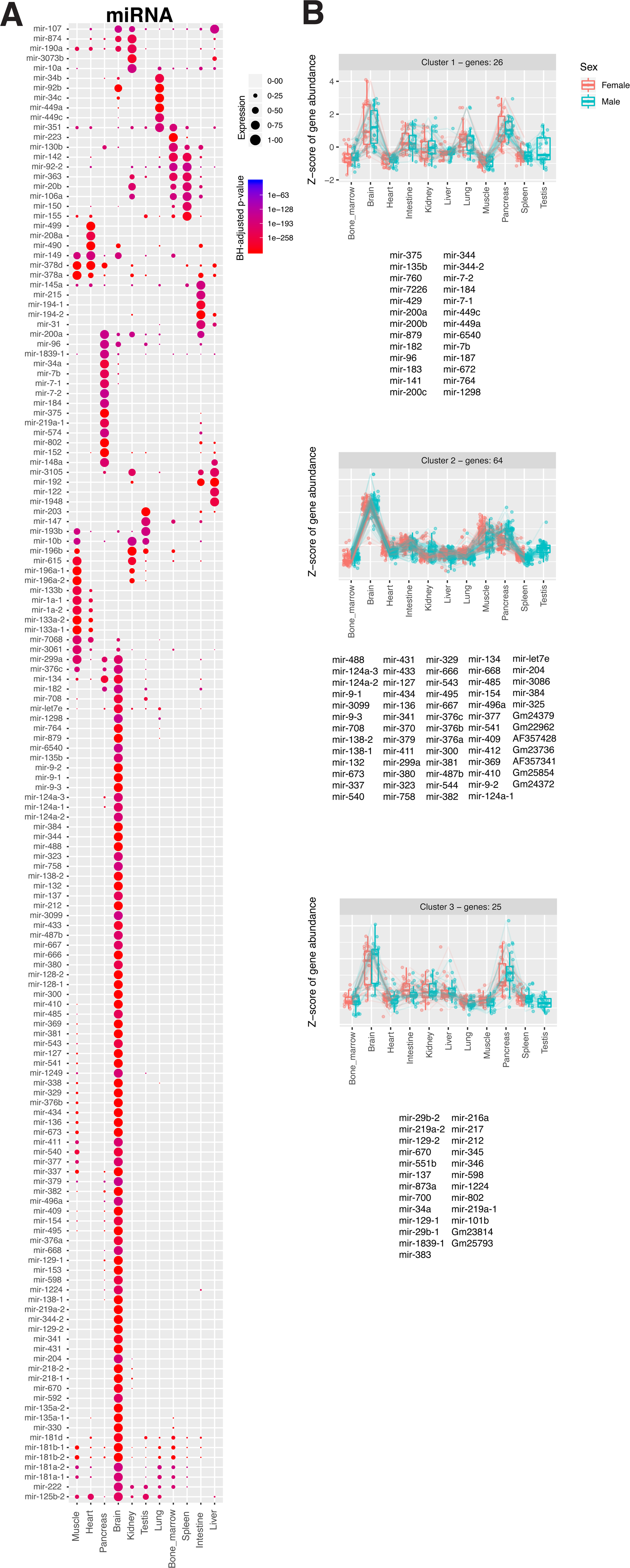
Tissue-specific patterns of sRNA expression. (A) Expression of top 150 tissue-specific GENCODE-annotated miRNA (BH-adjusted <0.01 in LRT test) across tissues. (B) Three gene clusters identified based on 250 top tissue-specific genes (lowest BH-adjusted and basemean>3).

**Figure S4.**
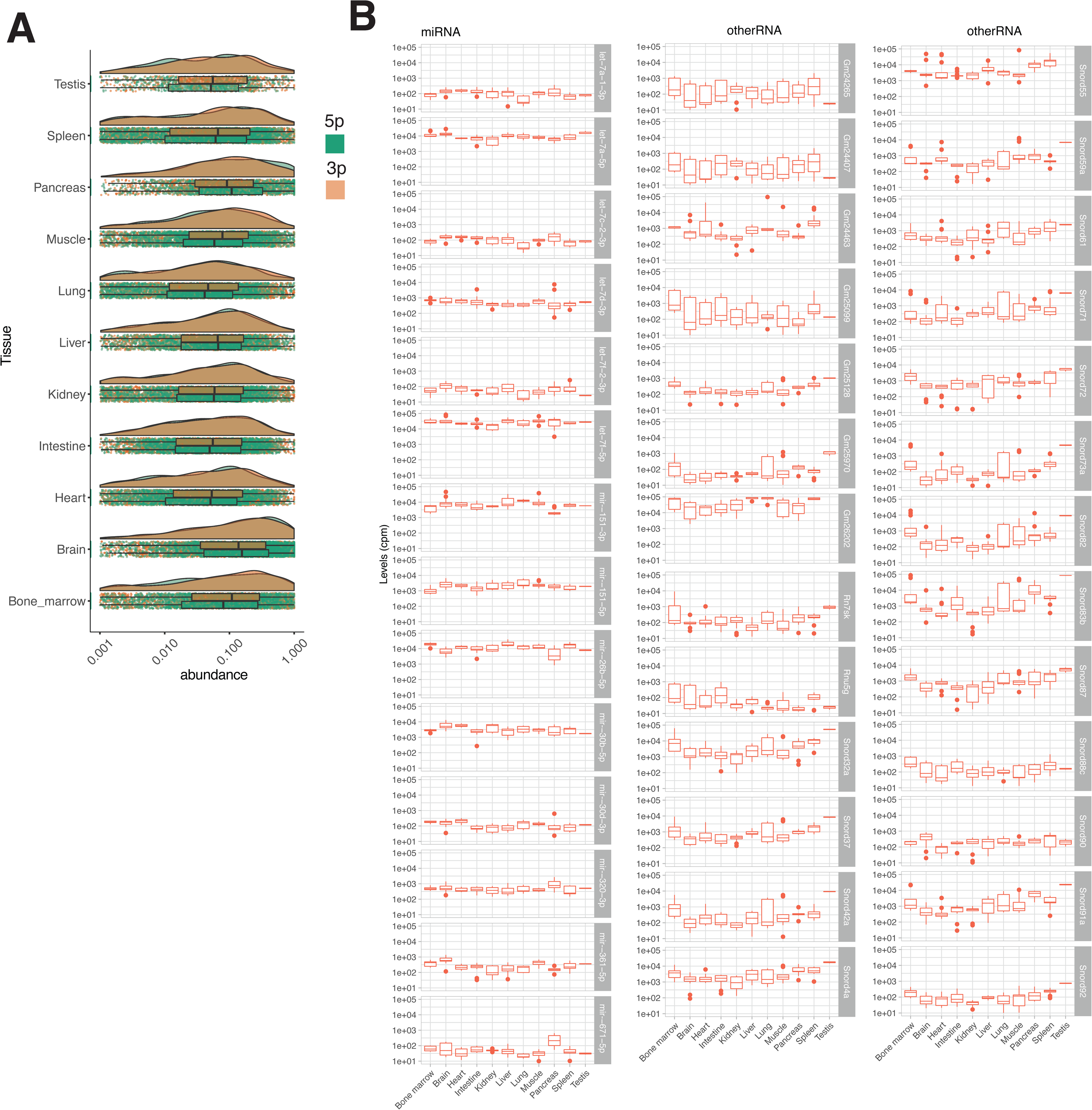
miRNA expression scores across eleven tissues. (A) Normalized abundance of 5p and 3p arms of all detected miRNAs in the profiles tissues. Each dot represents a mature miRNA. Density profiles were computed based on the levels of all 3p and 5p arms in each given tissue. (B) ncRNA ubiquitous across tissues.

**Figure S5.**
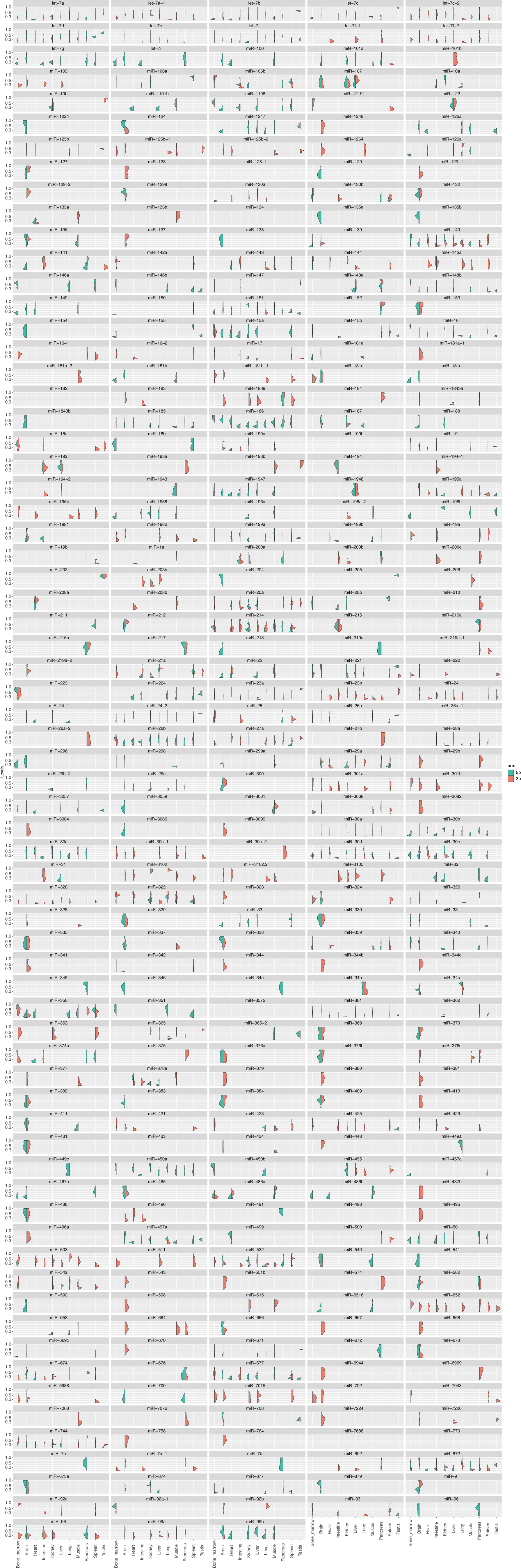
Levels of 5p and 3p arm of the same miRNA detected in different tissues. Y-axis represents normalized scaled counts.

**Figure S6.**
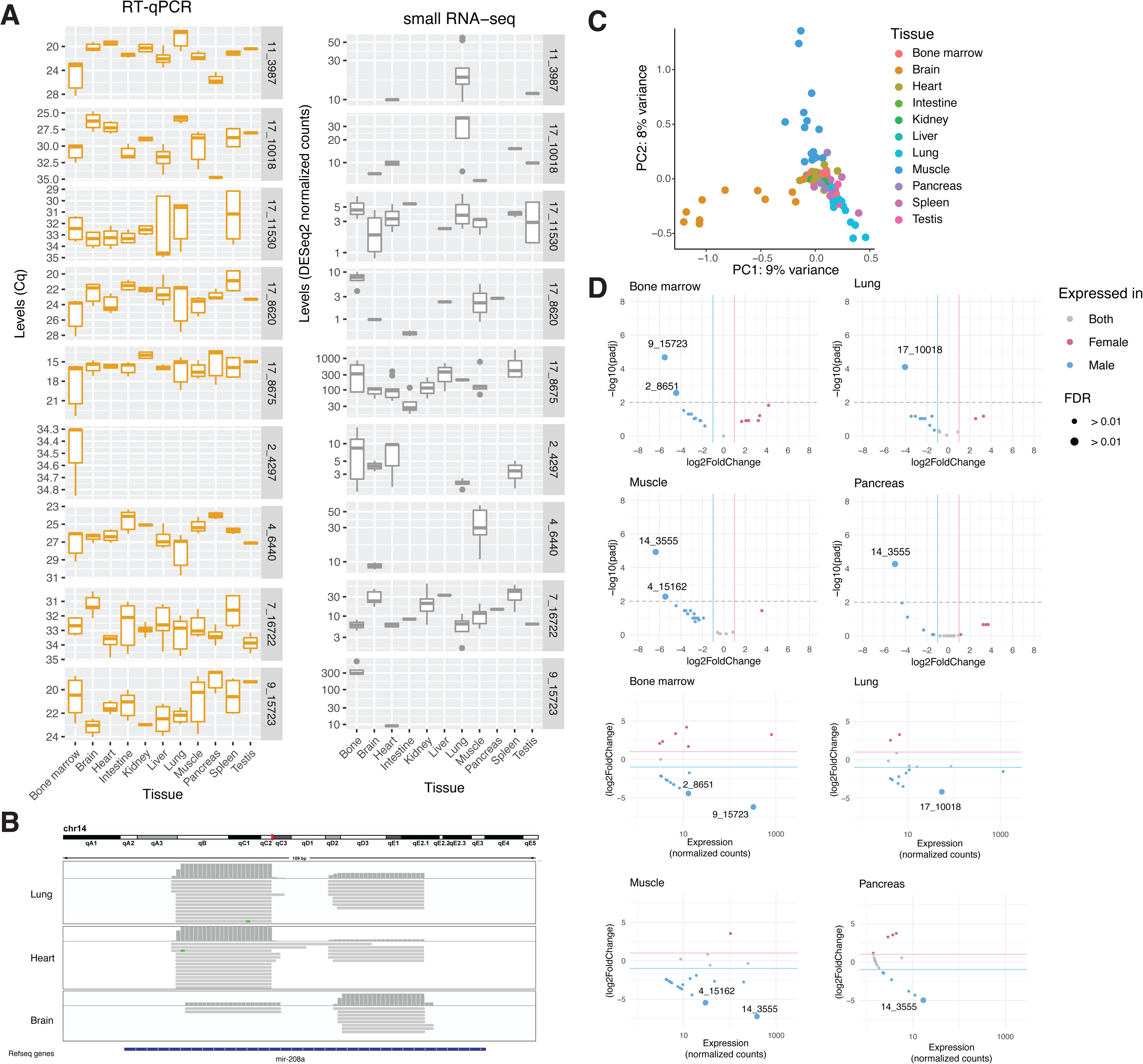
Novel miRNA. (A) Levels of 9 novel miRNAs measured with RT-qPCR and RNA-seq. (B) Example of a novel miRNA identified in the present study, that maps to the opposite strand of *mir-802a* in the brain. (C) Principal component analysis performed on novel miRNAs detected in one or multiple mouse tissues at > 50 counts. (D) Volcano plots showing novel miRNA differentially expressed between female and male bone marrow, lung, muscle and pancreas.

**Figure S7.**
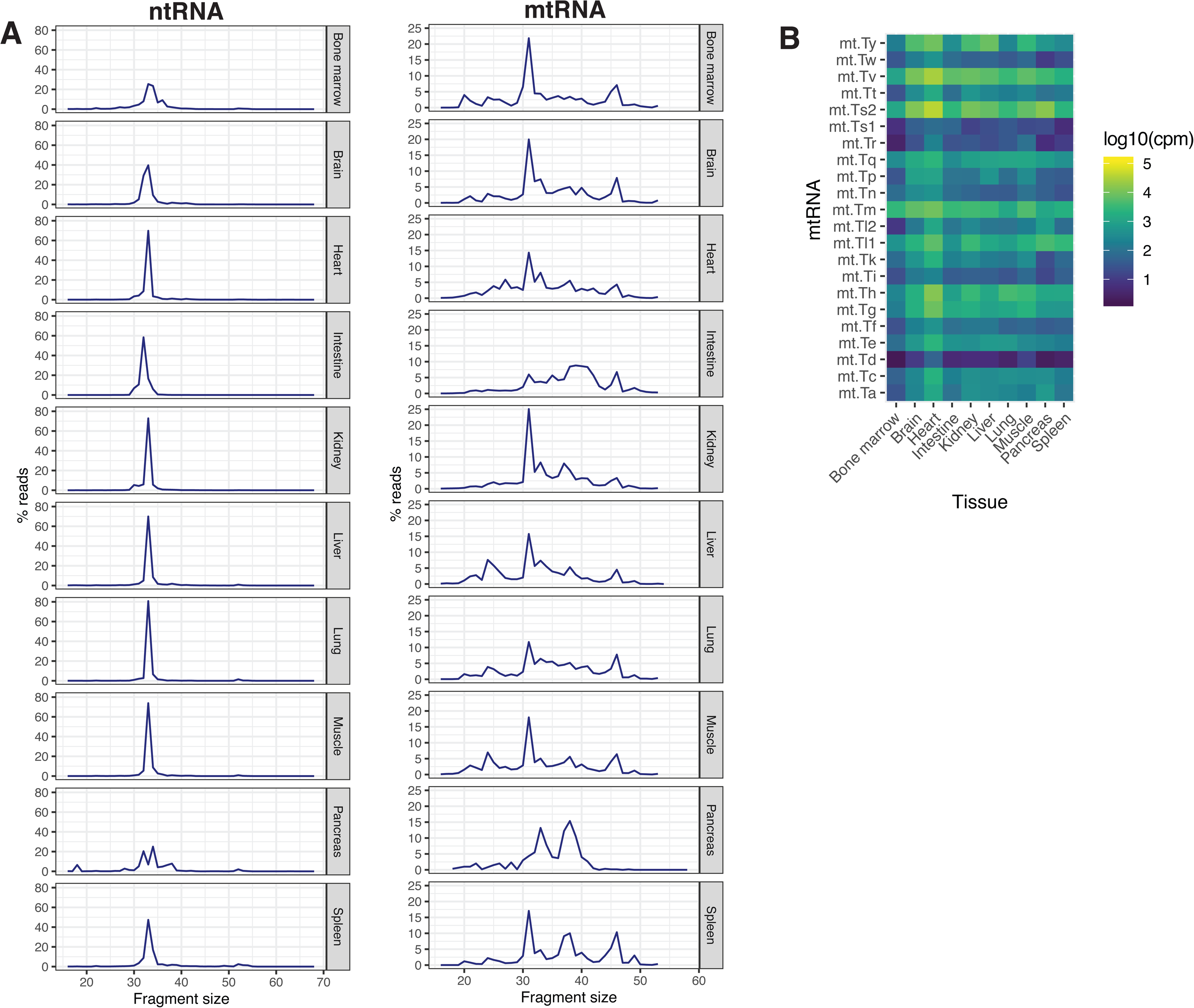
tRFs in mouse tissues. (A) Average length of either ntRNA or mtRNA fragments identified in each tissue. (B) Same as **Figure 4D** but for mtRNA fragments.

**Figure S8.**
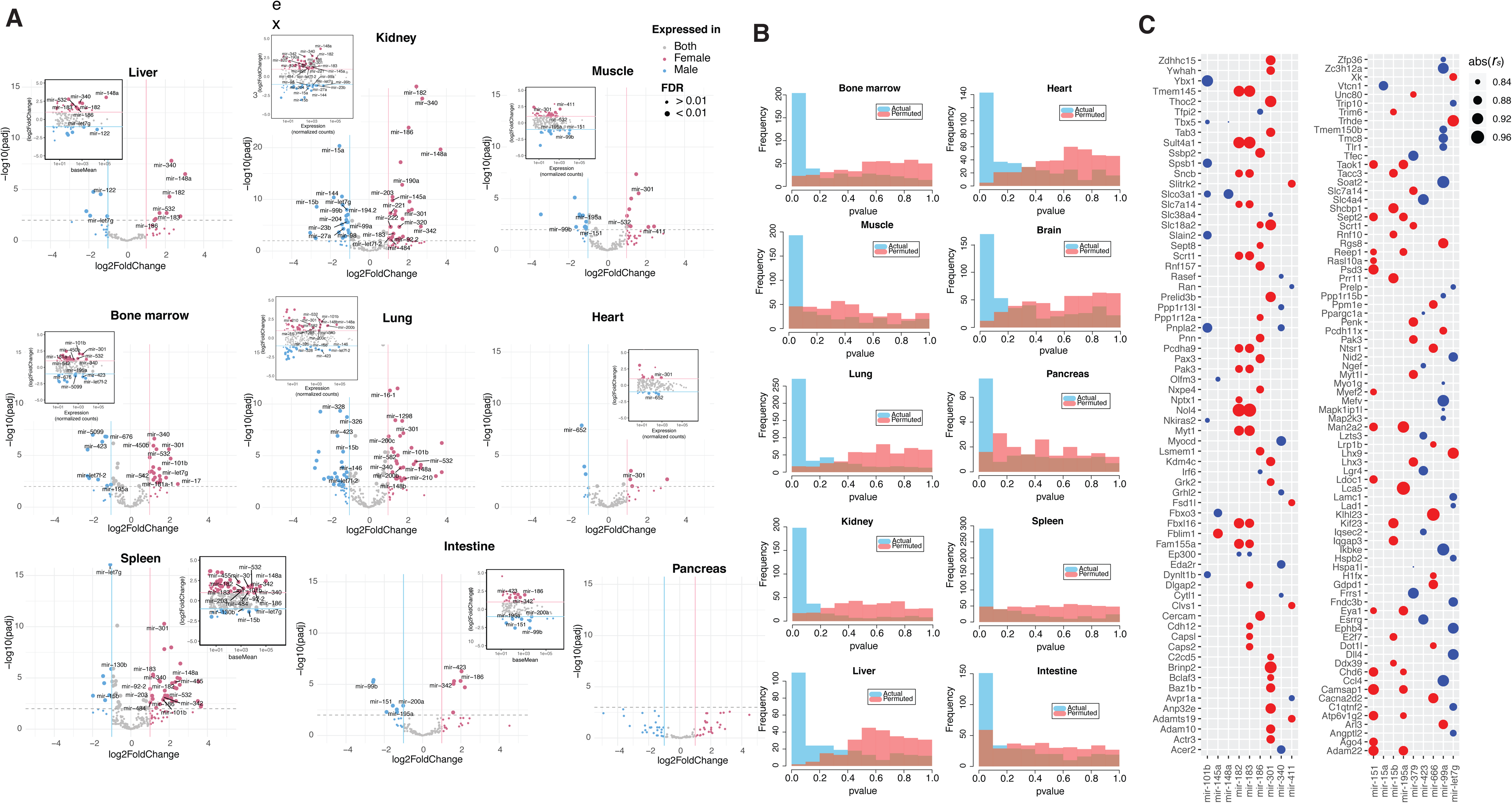
Sex-specific expression of miRBase-annotated miRNA. (A) Volcano plots showing annotated miRNAs differentially expressed between female and male intestine, spleen, liver and pancreas. (B) Differential expression analysis, NULL hypothesis testing. Distribution of *DESeq2*-derived p-values computed for miRNAs differentially expressed between actual and permuted female and male samples. (C) Spearman correlation of miRNA expression and the expression of respective targets. A list of target mRNAs was obtained from TargetScan, DIANA, miRanda, or mirDB databases (Agarwal et al., 2015; Griffiths-Jones et al., 2007; Paraskevopoulou et al., 2013; Wong and Wang, 2015). Only targets listed in two or more databases were used in this analysis. mRNA expression values for the respective tissues were retrived from the ENCODE database (Pennisi, 2012). Only *r_s_* scores of correlations that passed FDR<0.1 threshhold are shown. Positively correlated miRNA:mRNA scores are depicted in red, while negatively correlated – in blue.

**Figure S9.**
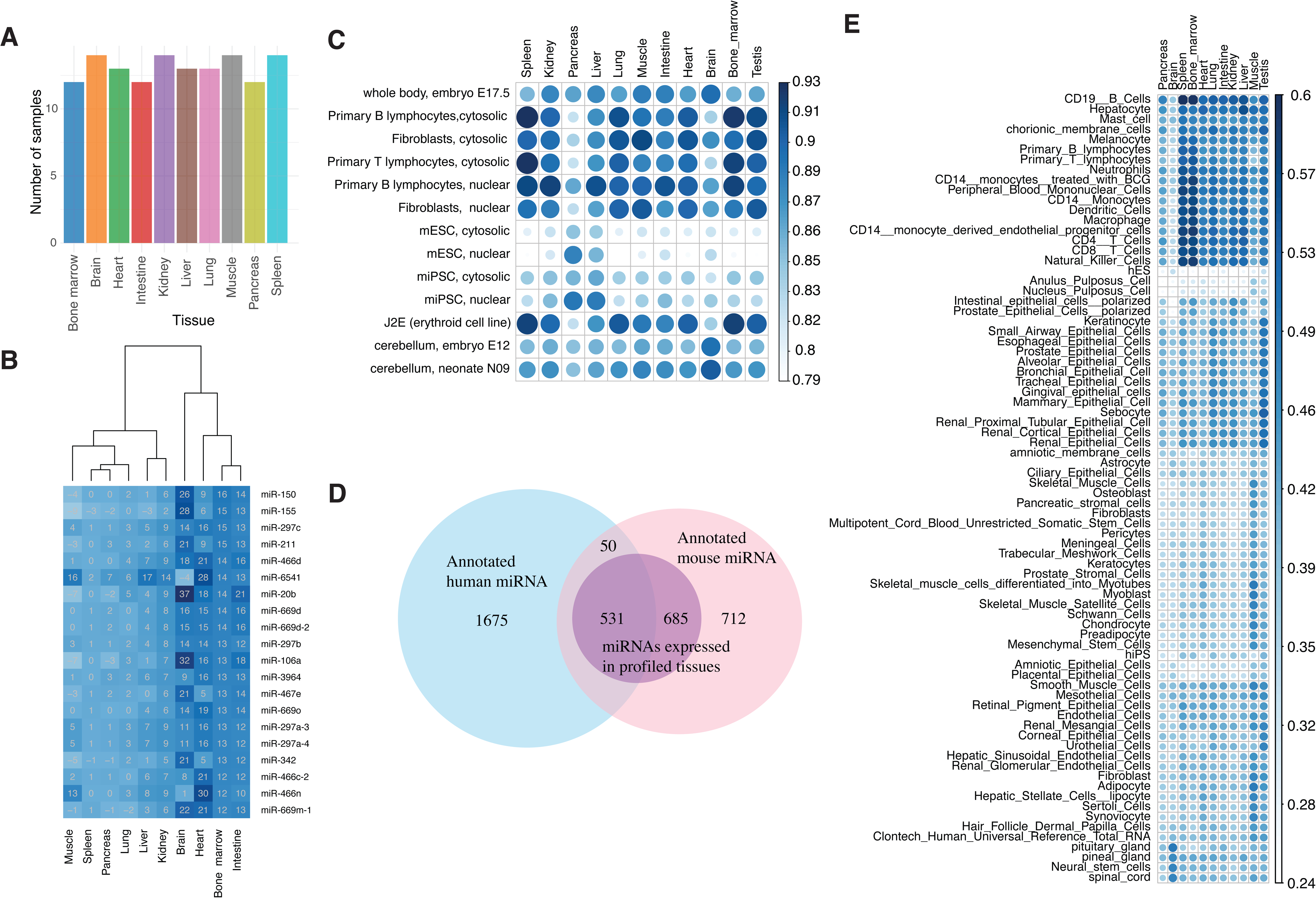
Comparison with previously published miRNA datasets generated for mammalian cell- and tissues-types. (A) RNA-seq samples used to train machine learning models. Left: number of train datasets corresponding to each tissue, right: examples of feature scores. (B) miRNAs assigned highest weights in defining bone marrow in SVM model. (C) Spearman correlation coefficients of comparison between expression scores obtained in this study and mouse miRNA-seq data generated by FANTOM5 (de Rie et al., 2017). (D) Number of miRNA orthologs used to compare FANTOM5 human and current mouse datasets. (E) same as A but for human FANTOM5 samples.

**Figure S10.**
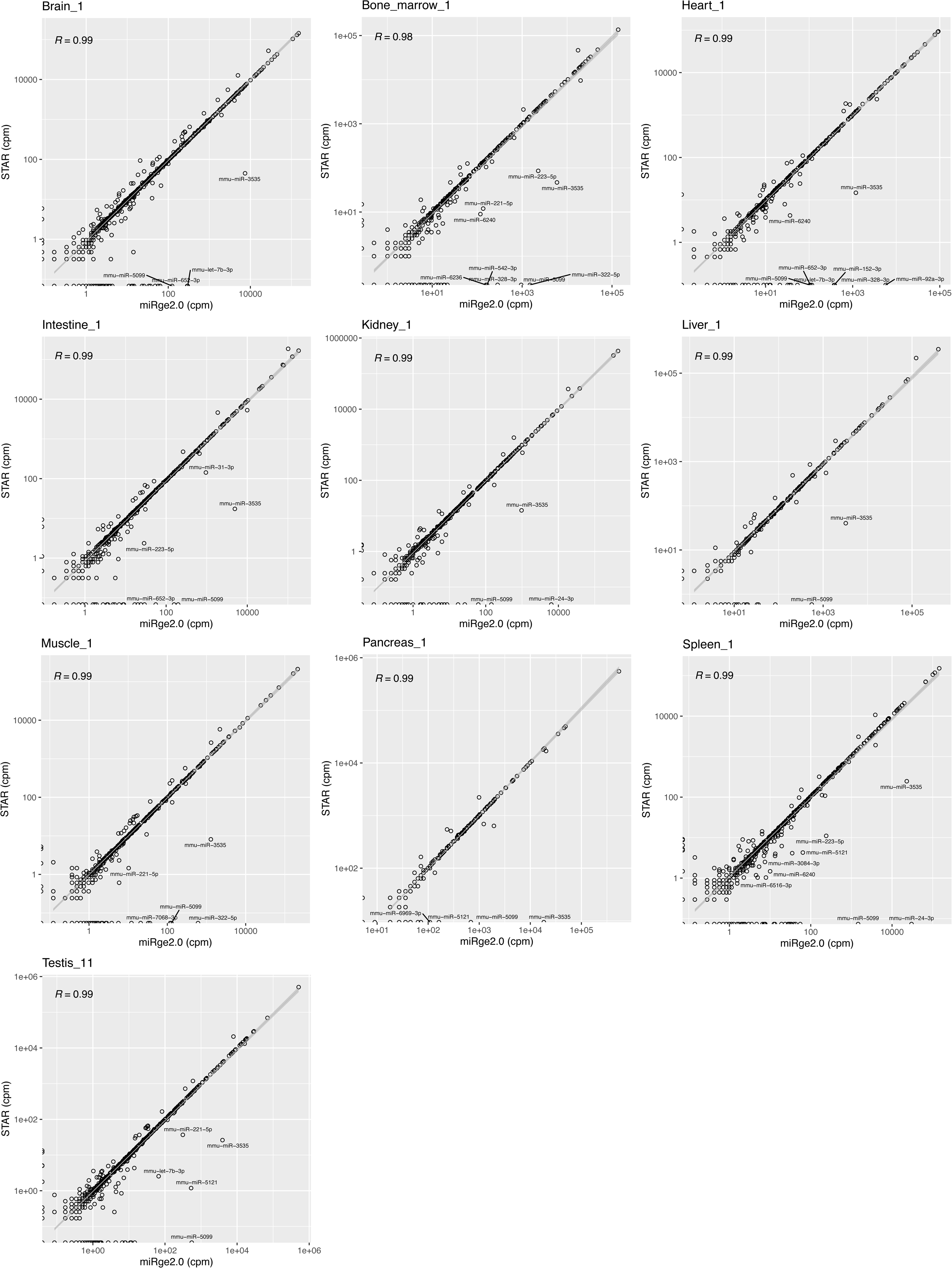
Comparison of mapping strategies. Correlation between miRNA levels obtained through mapping to the genome and derived from mapping to the nucleotide library (using miRge2.0 (Lu et al., 2018)).

## Supplementary tables

**Table S1.** Animals used in the study.

**Table S2.** Raw expression counts.

**Table S3.** Tissue-specificity scores.

**Table S4.** Novel miRNAs.

**Table S5.** ENCODE data used for ML predictions.

**Table S6.** Targets of sexually dimorphic miRNAs.

**Table S7.** ENCODE mRNA datasets used in the study.

